# Community Evaluation of Glycoproteomics Informatics Solutions Reveals High-Performance Search Strategies of Serum *N*- and *O*-Glycopeptide Data

**DOI:** 10.1101/2021.03.14.435332

**Authors:** Rebeca Kawahara, Anastasia Chernykh, Kathirvel Alagesan, Marshall Bern, Weiqian Cao, Robert J. Chalkley, Kai Cheng, Matthew S. Choo, Nathan Edwards, Radoslav Goldman, Marcus Hoffmann, Yingwei Hu, Yifan Huang, Jin Young Kim, Doron Kletter, Benoit Liquet-Weiland, Mingqi Liu, Yehia Mechref, Bo Meng, Sriram Neelamegham, Terry Nguyen-Khuong, Jonas Nilsson, Adam Pap, Gun Wook Park, Benjamin L. Parker, Cassandra L. Pegg, Josef M. Penninger, Toan K. Phung, Markus Pioch, Erdmann Rapp, Enes Sakalli, Miloslav Sanda, Benjamin L. Schulz, Nichollas E. Scott, Georgy Sofronov, Johannes Stadlmann, Sergey Y. Vakhrushev, Christina M. Woo, Hung-Yi Wu, Pengyuan Yang, Wantao Ying, Hui Zhang, Yong Zhang, Jingfu Zhao, Joseph Zaia, Stuart M. Haslam, Giuseppe Palmisano, Jong Shin Yoo, Göran Larson, Kai-Hooi Khoo, Katalin F. Medzihradszky, Daniel Kolarich, Nicolle H. Packer, Morten Thaysen-Andersen

## Abstract

Glycoproteome profiling (glycoproteomics) is a powerful yet analytically challenging research tool. The complex tandem mass spectra generated from glycopeptide mixtures require sophisticated analysis pipelines for structural determination. Diverse software aiding the process have appeared, but their relative performance remains untested. Conducted through the HUPO Human Proteome Project – Human Glycoproteomics Initiative, this community study, comprising both developers and users of glycoproteomics software, evaluates the performance of informatics solutions for system-wide glycopeptide analysis. Mass spectrometry-based glycoproteomics datasets from human serum were shared with all teams. The relative team performance for *N*- and *O*-glycopeptide data analysis was comprehensively established and validated through orthogonal performance tests. Excitingly, several high-performance glycoproteomics informatics solutions were identified. While the study illustrated that significant informatics challenges remain, as indicated by a high discordance between annotated glycopeptides, lists of high-confidence (consensus) glycopeptides were compiled from the standardised team reports. Deep analysis of the performance data revealed key performance-associated search variables and led to recommendations for improved “high coverage” and “high accuracy” glycoproteomics search strategies. This study concludes that diverse software for comprehensive glycopeptide data analysis exist, points to several high-performance search strategies, and specifies key variables that may guide future software developments and assist informatics decision-making in glycoproteomics.

## Introduction

Protein glycosylation, the attachment of complex carbohydrates (glycans) to discrete sites on proteins, plays diverse roles in biology^1^. Glycoproteomics, the system-wide analysis of intact glycopeptides, has evolved from proteomics and glycomics by the recognised benefits of concertedly studying glycan structures, modification sites and protein carriers at scale within a single experiment^2, 3^.

Facilitated by recent advances in separation science, mass spectrometry (MS) and informatics, glycoproteomics has matured over past decades and is now ready to tackle biological questions and generate new insights into the heterogeneous glycoproteome in biological systems^4–7^. While glycoproteomics studies now routinely report thousands of *N-* and *O*-glycopeptides^8^, accurate identification of glycopeptides from large volumes of mass spectral data remains a bottleneck. The annotation process of glycopeptide MS/MS data is highly error-prone due to the challenging task of correctly assigning both the glycan composition, modification site(s) and peptide carrier^9–11^. As a result, glycopeptides reported in glycoproteomics papers are frequently misidentified or suffer from ambiguous annotation even in studies attempting to control the false discovery rate (FDR) of assignments.

Diverse fragmentation modes including resonance-activation collision*-*induced dissociation (CID), beam-type CID (higher-energy collisional dissociation, HCD) and electron*-*transfer dissociation (ETD) have proven valuable for glycoproteomics^12–15^. When applied in concert, now possible, for example, on Orbitrap Tribrid mass spectrometers, these fragmentation strategies provide complementary structural information of glycopeptides. Briefly, HCD-MS/MS informs on the peptide carrier and produces useful diagnostic glycan fragments enabling glycopeptide classification and deduction of generic glycan compositions, ETD-MS/MS reveals in favourable cases the modification site and peptide identity, while resonance-activation CID-MS/MS informs primarily on the glycan composition, sequence and topology^16, 17^. Hybrid-type fragmentation strategies including ETciD and EThcD are becoming popular given their ability to generate information*-*rich glycopeptide spectra containing multiple fragment types^18^. Accurate mass measurements (<5-10 ppm) at high resolution of precursor and product ions available on most contemporary instruments are essential in glycoproteomics. Despite these exciting advances, unambiguous glycopeptide identification remains challenging. Informatics advances are therefore required to ensure accurate glycoproteome profiling to further the field^19^.

Glycoproteomics has experienced the development of diverse commercial and academic software showing promise for precise annotation and identification of glycopeptides from MS/MS data^20, 21^. While some of these tools are already well-established and widely applied in glycoproteomics^22^, the relative performance of software available to the community remain untested leaving a critical knowledge gap that hinders rapid progress in the field.

Facilitated by the HUPO Human Proteome Project – Human Glycoproteomics Initiative (HPP-HGI), we here perform a comprehensive community-based evaluation of existing informatics solutions for large-scale glycopeptide analysis. While informatics challenges undoubtedly still exist in glycoproteomics, our study highlights that several computational tools, some already demonstrating high performance, others considerable potential, are available to the community. Importantly, key performance-associated search parameters and high-performance search strategies were identified, findings that may aid software developers and users to improve the challenging glycoproteomics data analysis in the immediate future.

## Results

### Study design and overview

Two glycoproteomics data files (File A-B) of human serum were acquired (**Figure 1a**). A synthetic *N*-glycopeptide was included as a positive control. File A-B were generated using HCD-ETciD-CID-MS/MS and HCD-EThcD-CID-MS/MS, respectively, to accommodate most search engines. Serum is a well-characterised biospecimen displaying profound heterogeneity of *N-* and *O-*glycoproteins^23–25^. Thus, File A-B displayed characteristics (file size, complexity, type) similar to data typically encountered in glycoproteomics^26–29^.

**Figure 1.**
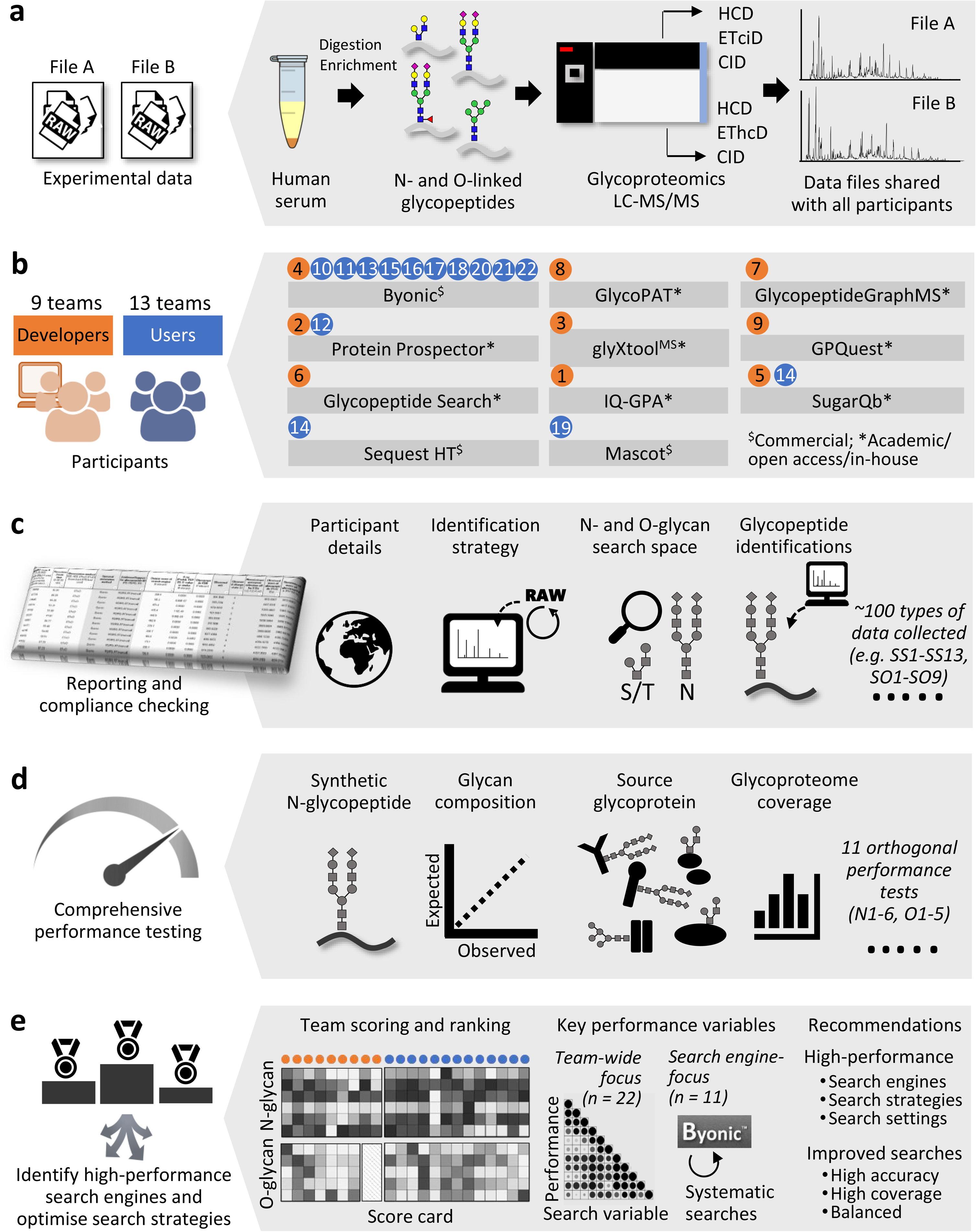
Study overview. **a**. Two glycoproteomics data files of human serum (File A-B) were generated and shared with participants. **b**. Participants comprising both developers (orange) and users (blue, team identifiers indicated) employed diverse search engines to complete the study. **c**. Teams returned a common reporting template capturing details of the applied search strategy including key search settings (SS1-SS13) and search output (SO1-SO9, **Table 1**) and identified glycopeptides. **d**. Complementary performance tests (N1-N6, O1-O5, **Table 1**) were used to comprehensively evaluate the ability of teams to identify *N-* and *O-*glycopeptides. **e**. The performance profiles were used to separately score and rank the developers and users. Diverse team-wide and search engine-centric (Byonic-focused) approaches were employed to identify performance-associated variables and high-performance search strategies.

**Table 1.** Overview of the symbols used in this study to refer to the key search settings (SS1-SS13), search output (SO1-SO9) and performance tests (N1-N6, O1-O5) applied to establish the relative team performance for glycopeptide data analysis. ^#^Average of output data reported by each team. AA, amino acid residues. While reported glycoPSMs were considered search output (SO9), unique glycopeptides (site-localisation not considered) were used to score the glycoproteome coverage (N4, O3).

| Search settings |  | Range or definition of category<br>(team count) |
| --- | --- | --- |
| SS1 | *N*-glycan search space | 23 - 381 unique glycan compositions |
| SS2 | *O*-glycan search space | 3 - 223 unique glycan compositions |
| SS3 | Search engine(s) applied | Byonic (11), Protein Prospector (2), GlycoPAT (1), GlycopeptideGraphMS (1), glyXtool<sup>MS</sup> (1), GPQuest (1), Other (5) |
| SS4 | Type of search engine | Academic/open access/in-house = 0 (7), Commercial = 1 (15) |
| SS5 | Spectral calibration post-acquisition | No = 0 (18), Yes = 1 (4) |
| SS6 | Protease specificity | Non-tryptic (*N*- or *O*-ragged or non-specific) = 0 (8), Tryptic = -1 (14) |
| SS7 | Missed peptide cleavages permitted | 0 - 2 missed cleavages |
| SS8 | Variable peptide modification(s) (non-glycan) | 0 - 14 non-glycan modification types |
| SS9 | Maximum glycans per peptide | 1 - 5 glycans/peptide |
| SS10 | Maximum other variable modifications | 0 - 5 variable modifications/peptide |
| SS11 | Precursor ion mass error permitted | Low (<5 ppm) = 0 (6), Medium (5-10 ppm) = 1 (14), High (>10 ppm) = 2 (2) |
| SS12 | Product ion mass error permitted | Low (<5 ppm) = 0 (0), Medium (5-10 ppm) = 1 (9), High (>10 ppm) = 2 (13) |
| SS13 | Peptide/protein FDR (decoy/contaminant database) | No decoy/contaminant = 0 (3), Only decoy or contaminant = 1 (9), Both decoy and contaminant = 2 (10) |
| Search output |  | Range<br>(*N*- or *O*-glycosylation) |
| SO1 | Glycopeptide LC retention time<sup>#</sup> | 41.7 - 57.2 min (NG)<br>26.2 - 55.8 min (OG) |
| SO2 | Glycopeptide *m/z* (observed)<sup>#</sup> | *m/z* 712.9 - 1199.4 (NG)<br>*m/z* 619.6 - 1229.0 (OG) |
| SO3 | Glycopeptide charge state (observed)<sup>#</sup> | *z* = 3.9 - 4.3 (NG)<br>*z* = 3.1 - 5.4 (OG) |
| SO4 | Monoisotopic correction (off-by-X, positive values)<sup>#</sup> | 0 - 2.1 Da (NG)<br>0 - 2.0 Da (OG) |
| SO5 | Glycopeptide mass ([M+H], observed)<sup>#</sup> | 3144.6 - 4913.0 Da (NG)<br>1892.9 - 6057.1 Da (OG) |
| SO6 | Glycopeptide actual mass error (observed, positive values)<sup>#</sup> | 0.5 - 2.8 ppm (NG)<br>1.1 - 5.9 ppm (OG) |
| SO7 | Glycopeptide length<sup>#</sup> | 16.9 - 26.8 AA (NG)<br>14.5 - 38.9 AA (OG) |
| SO8 | Glycan mass (calculated from reported glycopeptides)<sup>#</sup> | 1880.8 - 2410.1 Da (NG)<br>195.8 - 2216.8 Da (OG) |
| SO9 | Reported GlycoPSMs | 49 - 2122 glycoPSMs (NG)<br>5 - 578 glycoPSMs (OG) |
| **Performance tests** |  | **Description of scoring method** |
| N1 | Synthetic *N*-glycopeptide | Identification accuracy (specificity, %) multiplied by coverage (sensitivity, %) of a synthetic *N*-glycopeptide (**Supplementary Table 5**) |
| N2 | *N*-glycan composition | Pearson correlation ($R^2$) between the expected<sup>23</sup> and observed *N*-glycan distribution in human serum (**Supplementary Table 6**) |
| N3 | Source *N*-glycoprotein | Specificity and sensitivity of reported source glycoproteins relative to expected serum glycoproteins<sup>23, 39</sup> (**Supplementary Table 7**) |
| N4 | *N*-glycoproteome coverage | Unique *N*-glycopeptides (unique peptide sequence and *N*-glycan composition) (**Supplementary Table 8**) |
| N5 | Commonly reported ‘consensus’ *N*-glycopeptides | Proportion of reported glycopeptides of the consensus glycopeptides commonly reported by >50% teams (**Supplementary Table 9**) |
| N6 | NeuGc and multi-Fuc *N*-glycopeptides (absence) | Proportion of reported glycoPSMs <u>not</u> containing NeuGc and multi-Fuc of all reported glycoPSMs. Only applicable if NeuGc/Fuc $\geq 2$ glycans were included in glycan search space (**Supplementary Table 10**) |
| O1 | *O*-glycan composition | Pearson correlation ($R^2$) between the expected<sup>40</sup> and observed *O*-glycan distribution in human serum (**Supplementary Table 11**) |
| O2 | Source *O*-glycoprotein | Specificity and sensitivity of reported source glycoproteins relative to expected serum glycoproteins<sup>4, 41, 42</sup> (**Supplementary Table 12**) |
| O3 | *O*-glycoproteome coverage | Unique *O*-glycopeptides (unique peptide sequence and *O*-glycan composition) (**Supplementary Table 13**) |
| O4 | Commonly reported ‘consensus’ *O*-glycopeptides | Proportion of reported glycopeptides of the consensus *O*-glycopeptides commonly reported by >30% teams (**Supplementary Table 14**) |
| O5 | NeuGc and multi-Fuc *O*-glycopeptides (absence) | Proportion of reported glycoPSMs <u>not</u> containing NeuGc and multi-Fuc containing glycoPSMs of all reported glycoPSMs. Only applicable if NeuGc/Fuc $\geq 2$ glycans were included in glycan search space (**Supplementary Table 15**) |

File A-B were shared with all 22 participating teams who classified themselves as developers (9 teams) or users (13 teams) of glycoproteomics software (**Figure 1b**, **Extended Data Figure 1a-b**). All teams identified *N-* and *O-*glycopeptides from File A-B and reported their approaches and identifications in a standardised reporting template. Most developers (five teams) and users (eight teams) were experienced in glycoproteomics (>10 years). Participants were from North and South America, Europe, Asia, and Oceania, (**Extended Data Figure 1c-d**).

Only File B was processed by all teams (**Extended Data Figure. 1e**). While most participants reported on spectra acquired with multiple fragmentation methods, a few teams used only HCD- or EThcD-MS/MS for the identifications (**Extended Data Figure 1f-g**).

Participants used diverse search engines (**Figure 1b**, **Extended Data Figure 1h**). Some search engines were used as stand-alone tools or with other software while others were applied with pre-/post-processing tools to aid the identification (**Extended Data Figure 1i**). The developers used 9 different glycopeptide-centric search engines including Team 1: IQ-GPA v2.5^30^, Team 2: Protein Prospector v5.20.23^31^, Team 3: glyXtool^MS^ v0.1.4^32^, Team 4: Byonic v2.16.16^33^, Team 5: Sugar Qb^34^, Team 6: Glycopeptide Search v2.0alpha^35^, Team 7: GlycopeptideGraphMS v1.0^36^/Byonic^33^, Team 8: GlycoPAT v2.0^37^, Team 9: GPQuest v2.0^38^. Amongst the 13 users, 10 teams (∼75%) used Byonic (Team 10, 11, 13, 15-18, 20-22), while only few teams used Protein Prospector (Team 12), SugarQB/Sequest HT (Team 14), and Mascot (Team 19) (**Supplementary Table 1**).

File A/B contained 8,737/9,776 HCD-MS/MS scans of which 5,485/6,148 (∼63%) spectra contained glycopeptide-specific oxonium ions (e.g. *m/z* 204.0867) used for ETciD-/EThcD-/CID-MS/MS triggering (**Extended Data Figure 2a-b**). Amongst all potential glycopeptide MS/MS spectra (File A/B, 16,445/18,444, considering all fragmentation modes), 3,402/4,982 (20.7%/27.0%) non-redundant (unique) glycopeptide-to-spectrum matches (glycoPSMs) were collectively reported by participants. Most teams reported on HCD- and EThcD-MS/MS data, while only few teams used CID- and ETciD-MS/MS. Similar charge distribution (most frequently quadruply-charged precursors) was observed for glycopeptides reported from different fragmentation modes (**Extended Data Figure 2c-d**).

A wealth of data was collected via a comprehensive reporting template. The team reports covered intricate details of the employed search strategies and identified glycopeptides (**Figure 1c**). Details of the applied search settings were captured including permitted peptide modifications, mass tolerance, post-search filtering criteria (**Supplementary Table 1**), and glycan search space (**Supplementary Table 2**). The search settings (SS1-SS13, **Table 1**) varied considerably across teams.

Additionally, diverse output data arising from the glycopeptide identifications were captured (**Supplementary Table 3**). The output data also varied across teams (**Supplementary Table 4**). Analysis of key search output variables (SO1-SO9, **Table 1**) revealed that the reported *N-* and *O-*glycopeptides, as expected, showed different characteristics (e.g. LC retention time, glycan mass) while other characteristics (e.g. observed precursor *m/z*) were similar between the two analyte classes (**Extended Data Figure 3**). Analysis of SO1-SO9 data also demonstrated that some teams reported highly discrepant output. For example, and without being able to link these observations to performance, the developers of Glycopeptide Search (Team 6) and GlycopeptideGraphMS (Team 7) reported glycopeptides with unusually low (z

= ∼3+) and high (∼5.5+) charge states relative to other teams (∼4.5+) (**Extended Data Figure 3c**). These output data comparisons may be valuable for developers to better understand, further develop, and ultimately improve their software.

The team performance was assessed using orthogonal performance tests that served to comprehensively evaluate the glycopeptide identification accuracy (specificity) and glycoproteome coverage (sensitivity), two key performance characteristics in glycoproteomics (**Figure 1d**). Six (N1-N6) and five (O1-O5) performance tests were carefully designed to assess the relative performance for *N*- and *O*-glycopeptide data analysis across teams (**Table 1**, **Supplementary Table 5-15**). Firstly, the ability to detect the synthetic *N*-glycopeptide in the datasets was assessed (N1). Further, the glycan compositions (N2, O1) and source glycoproteins (N3, O2) of reported glycopeptides were compared to the established serum glycome and against known serum glycoproteins^4, 23, 39–42^. To validate the use of literature to score teams, we performed manual site-specific glycoprofiling of four serum glycoproteins i.e. alpha-1-antitrypsin (A1AT), ceruloplasmin (CP), haptoglobin (HP) and immunoglobulin G1 (IgG1) and showed an excellent agreement (R^2^ = 0.85-0.99) with relevant literature of healthy human serum^43–46^ (**Extended Data Figure 4**). The glycoproteome coverage was simply the reported non-redundant glycopeptides (N4, O3). Finally, the ability to identify glycopeptides commonly reported by most teams (“consensus glycopeptides”) (N5, O4) and free of NeuGc and multi-Fuc features (N6, O5) was also scored. We ensured that NeuGc and multi-Fuc glycopeptides, unexpected glycofeatures in human serum^23, 47–49^, were indeed absent or rarely detected in File A-B (discussed below) allowing these to be deemed putative false positives for the purpose of scoring teams (**Extended Data Figure 5**).

The performance tests were used to score and rank teams (**Figure 1e**, **Supplementary Table 16**). The developer and user groups were not compared since they received different study instructions. The team scoring was validated using an independent glycoprotein-centric site-specific profiling test **(Supplementary Table 17**). Finally, performance data from both team-wide and search engine-centric approaches revealed performance-associated search variables and led to improved glycoproteomics search strategies (**Supplementary Table 18-19**).

### Overview of the reported glycopeptides

The below analyses were carried out using data reported from File B processed by all teams. The total *N-*glycoPSMs (49-2,122) and source glycoproteins (9-168) reported by the 22 teams varied dramatically (**Figure 2a**, **Supplementary Table 3**). In line with literature of human serum *N*-glycosylation^23, 24^, the reported *N-*glycopeptides carried mainly complex-type *N-* glycans (92.6%, average across teams) while relatively few oligomannosidic (6.4%) and truncated (Hex_<4_HexNAc_<4_) (1.0%) *N-*glycopeptides were reported. The applied *N-*glycan search space spanned an equally wide range (23-381 compositions) comprising mostly complex-type *N*-glycans (89.1%) and the less heterogeneous oligomannosidic (5.9%) and truncated (5.0%) *N-*glycans. No associations were found between the size of the *N-*glycan search space and reported *N-*glycoPSM counts (Pearson R^2^ = 0.115). Unexpected glycan compositions including NeuGc- and multi-Fuc-containing complex-type *N-*glycans, which are negligible features of human serum glycoproteins^23, 47–49^, were not only included in the glycan search space (up to 26.5% and 28.9%, respectively), but also reported (up to 20.6% and 5.0%) by some teams. The absence of NeuGc and rarity of multi-Fuc glycopeptides in the shared data was supported by a lack of diagnostic fragment ions for NeuGc (*m/z* 290/308), scarcity of antenna Fuc ions (*m/z* 512/803) and the frequent mis-annotation of MS/MS spectra claimed to correspond to NeuGc and multi-Fuc glycopeptides (**Extended Data Figure 5**). While only infrequently detected, multi-Fuc glycopeptides were, however, evidently present in our data as supported by manual spectral annotation (**Extended Data Figure 5d**).

**Figure 2.**
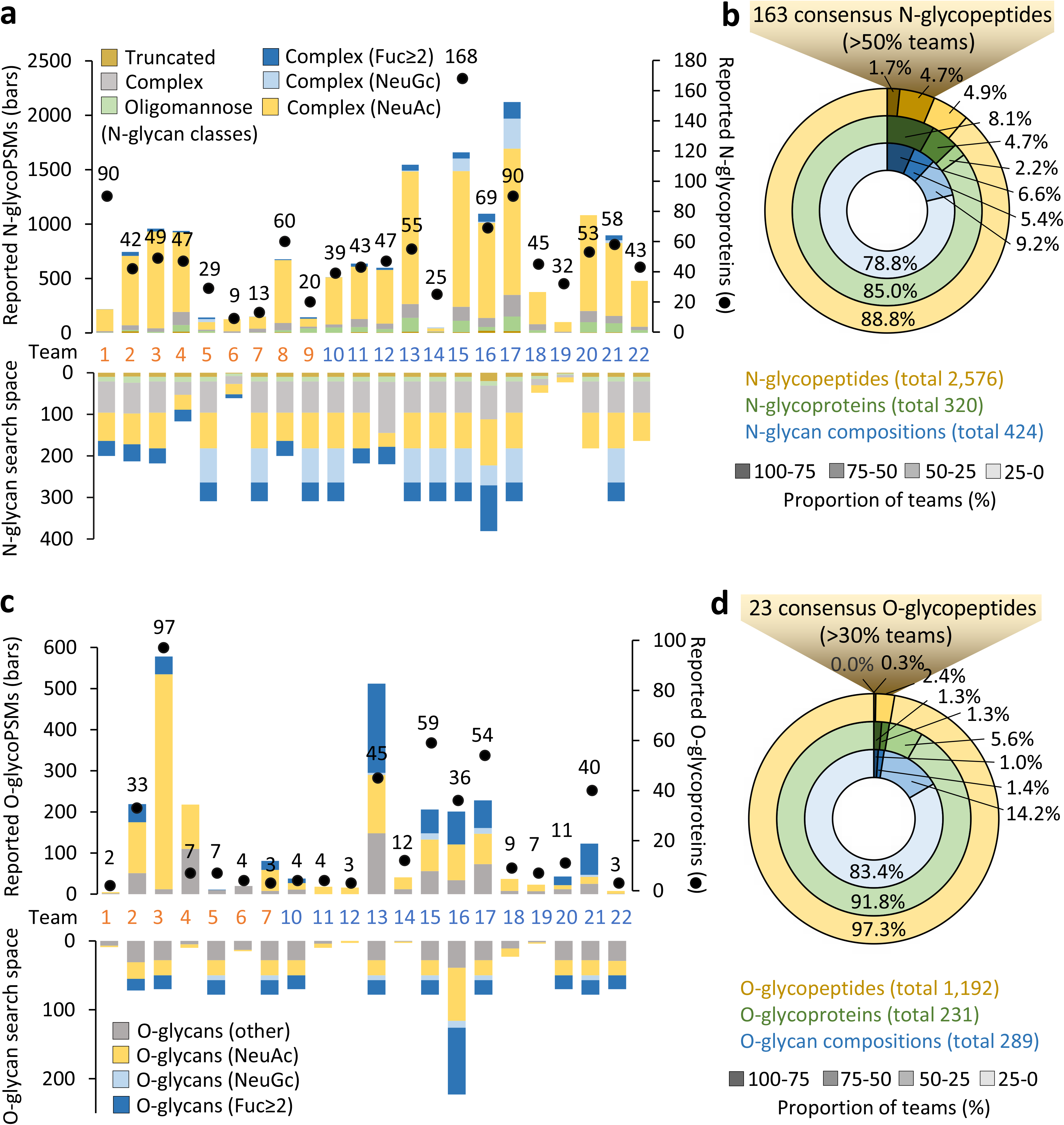
Overview of the glycopeptides reported across teams. Team 8-9 did not perform *O*-glycopeptide analysis. **a,c.** Distribution of the reported glycoPSMs (bars), unique source glycoproteins (dots) and applied glycan search space (mirror, see also **Supplementary Table 2-3**) across teams. **b,d.** Proportion of glycopeptides, source glycoproteins and glycan compositions commonly reported by teams. The high-confidence ‘consensus’ glycopeptides have been made publicly available (GlyConnect Reference ID 2943).

Collectively, 2,556 unique *N-*glycopeptides (defined herein as unique peptide sequences and glycan compositions), covering 320 different source *N-*glycoproteins and 424 different *N-* glycan compositions were reported across teams (**Figure 2b**, **Supplementary Table 6-7, 9**). Of these, only 43 *N*-glycopeptides (1.7%), 26 source *N*-glycoproteins (8.1%) and 28 *N*-glycan compositions (6.6%) were commonly reported by at least 75% of teams (see **Extended Data Figure 6a** for an example of congruent spectral annotation across teams). Most glycopeptides, however, were commonly reported by only few teams likely due to frequent mis-annotation of the spectral data (**Extended Data Figure 6b**).

Significantly fewer, but equally discrepant, *O-*glycopeptides (5-578 *O-*glycoPSMs) were reported by participants (**Figure 2c**, **Extended Data Figure 3i**). As expected, most reported *O-*glycopeptides carried Hex_1_HexNAc_1_NeuAc_1-2_^40, 50^. The applied *O*-glycan search space also varied dramatically (3-223 glycan compositions). Similar to the *N*-glycopeptide analysis, no association was observed between the *O-*glycan search space and reported *O-*glycoPSM counts (Pearson R^2^ = 0.118). Instead, many other associations were identified (discussed below). While seven teams included NeuGc in the applied *O*-glycan search space (up to 9.0%), only four teams reported NeuGc *O*-glycopeptides (up to 7.3%). In addition, 12 teams included multi-fucosylated glycans in the *O*-glycan search space (up to 43.5%); 11 of those teams reported multi-fucosylated *O*-glycopeptides (average of 28.6%, up to 61.8%). Both NeuGc and multi-fucosylated *O*-glycans are negligible features of human serum *O*-glycoproteins as supported by literature^40, 50^ and our own analyses (see above). The reported multi-fucosylated *O*-glycan compositions could, in principle, in some cases arise from multiple discrete *O*-glycans residing on the same peptide. Since *O*-glycosylation sites were inconsistently and/or ambiguously reported by most teams (see below) we were not able to assess this aspect further.

Collectively, 1,192 unique *O-*glycopeptides covering 231 different source *O-*glycoproteins and 288 different *O-*glycan compositions were identified, but surprisingly few *O-* glycopeptides were commonly reported across teams. Only 3 *O-*glycopeptides (0.3%), 6 source *O-*glycoproteins (2.6%) and 7 *O-*glycan compositions (2.4%) were commonly reported by at least half the teams (**Figure 2c**). Most *O*-glycopeptides were reported by a single or few teams.

Despite the discrepant reporting, high-confidence lists spanning 163 *N*- and 23 *O*-glycopeptides commonly reported by teams could be generated. Importantly, these consensus glycopeptides mapped to expected serum glycoproteins e.g. α-2-macroglobulin (UniProtKB, P01023) and haptoglobin (P00738) and carried expected serum *N*-glycans e.g. Hex_5_HexNAc_4_Fuc_0-1_NeuAc_2_ (GlyTouCan IDs, G09675DY/G22754FQ) and *O*-glycans e.g. Hex_1_HexNAc_1_NeuAc_1-2_ (G65285QO/G84906ML) that were biosynthetically related (**Extended Data Figure 7**), devoid of NeuGc and poor in multi-Fuc further supporting their correct identification. These high-confidence glycopeptides form an important reference to future studies of the human serum glycoproteome and have therefore been made publicly available (GlyConnect Reference ID 2943).

### High-performance informatics solutions for N-glycoproteomics

The relative team performance for *N*-glycoproteomics was comprehensively assessed using six independent performance tests (N1-N6) (**Table 1**, **Supplementary Table 5-10**). Amongst these performance tests, N1 scored the ability to accurately identify a synthetic *N*-glycopeptide in the sample (**Extended Data Figure 8**). Similar to the other performance tests, N1 was used to establish the relative team performance. Founded on a “ground truth”, the N1 data including the 12 manually annotated spectra all corresponding to the synthetic *N*-glycopeptide are particularly informative and may aid developers train algorithms and improve software to better annotate *N*-glycopeptide spectral data. The N1 data also supported observations made across the entire dataset (**Exten**ded Data Figure 2) confirming that glycopeptides were preferentially identified in charge state 4+ using HCD- and EThcD- MS/MS even when high-quality MS/MS data from other charge states and fragmentation modes were available.

In line with literature^2, 8^, most teams employed HCD- and/or EThcD-MS/MS for glycopeptide identification. While these two fragmentation modes displayed similar performance in tests scoring the glycan composition (N2, O1) and glycoproteome coverage (N4, O3), higher scores were achieved for EThcD-relative to HCD-based identifications in the source glycoprotein tests (N3, O2) (**Extended Data Figure 9**). Importantly, accurate glycosylation site localisation, not tested with this study (discussed below), is a recognised strength of EThcD-MS/MS data^5, 12^.

The performance tests were used to score and rank developers and users (**Figure 3a**). At a glance, the scorecard pointed to considerable team-to-team variations in the performance profiles suggesting that the applied software and search strategies exhibit markedly different strengths and weaknesses for *N*-glycoproteomics. As an example, IQ-GPA (Team 1) and GlycoPAT (Team 8) performed well (relative to other developers) in the *N-*glycan composition test (N2), while Protein Prospector (Team 2) and Byonic (Team 4) performed well in tests scoring the source *N-*glycoproteins (N3) and *N-*glycoproteome coverage (N4).

**Figure 3.**
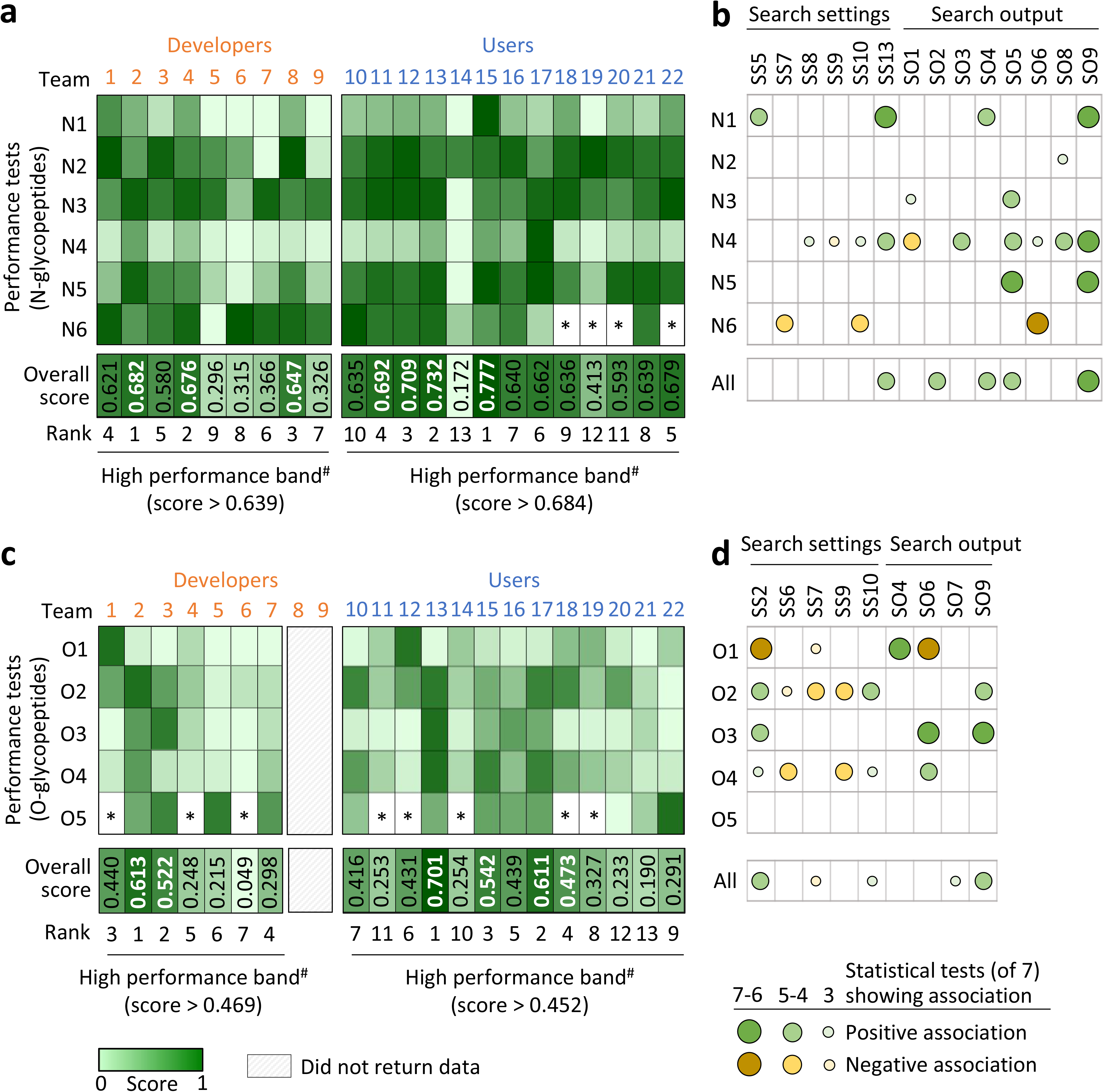
Team scoring/ranking and identification of performance-associated variables. **a,c.** Heatmap representation of normalised scores (range 0-1) from the glycopeptide performance tests (N1-N6, O1-O5, **Table 1**). Team 8-9 did not return *O*-glycopeptide data. See **Supplementary Table 5-16** for performance data. ^#^The top third performing teams (white bold) were placed in a high-performance band. Team scoring was validated using an independent assessment method (**Extended Data Figure 10**). *Performance could not be determined. **b,d.** Many study variables (search settings and search output) showed associations (negative or positive) with performance. See **Table 1** for variables. See **Supplementary Table 18** for statistics.

Overall, Protein Prospector (Team 2, overall score 0.682), Byonic (Team 4, 0.676) and GlycoPAT (Team 8, 0.647) were found to be high-performance software solutions for *N*-glycoproteomics. Notably, our scoring did not separate these three developers by any significant margin, but their overall performance was slightly higher than IQ-GPA (Team 1, 0.621) and glyXtool^MS^ (Team 3, 0.580) and substantially higher than other software (score range 0.296-0.366).

Supporting our scoring method, an independent assessment method based on the match between reported and actual site-specific *N*-glycoforms of four serum glycoproteins (A1AT, CP, HP, IgG1, thus founded on a “ground truth”) recapitulated the scoring profile across teams (R^2^ = 0.82) (**Extended Data Figure 10**). Further supporting the top ranking of Byonic and Protein Prospector, the best performing user teams employed Byonic (Team 11, 13, 15, score range 0.687-0.777) and Protein Prospector (Team 12, 0.709). Their overall performance scores were marginally higher than seven other Byonic users (Team 10, 16-18, 20-22, 0.593-0.679), but markedly higher than teams using SugarQb (Team 14, 0.172) and Mascot (Team 19, 0.413). Despite the similar overall performance amongst most user teams, not least the 10 Byonic users, their performance profiles differed markedly across the six performance tests.

We then explored the scorecard for software-independent performance-associated variables including the search settings (SS1-SS13) and search output (SO1-9) using seven different statistical methods (**Figure 3b**). Many statistically strong relationships were found revealing key performance-associated variables that either positively or negatively correlated with the glycopeptide identification efficiency. As an example, the use of decoy/contaminant databases (SS13) showed associations with performance in the synthetic *N*-glycopeptide test (N1) and high *N-*glycoproteome coverage (N4). Search strategies that allowed for a relatively high diversity and number of non-glycan variable peptide modifications (SS8, SS10) and few glycans per peptide (SS9) were also associated with high *N-*glycoproteome coverage (N4). As expected, allowing multiple missed peptide cleavages (SS7) and variable non-glycan modifications (SS10) in the search strategy correlated with higher glycopeptide FDRs as indicated by higher rates of NeuGc and multi-Fuc identifications (low N6 scores) (**Supplementary Table 18**).

The association analyses also identified many interesting relationships between the search output and performance (**Figure 3b**). Intuitively, teams that reported many *N*-glycoPSMs (SO9) performed well in the synthetic *N-*glycopeptide test (N1), had a higher *N-* glycoproteome coverage (N4) and identified more consensus *N-*glycopeptides (N5). Further, teams that reported glycopeptides featuring a relatively high glycan mass (SO8) more often identified the correct glycan composition (N2), while teams that reported glycopeptides exhibiting relatively high molecular masses (SO5) more often identified the correct source *N-* glycoproteins (N3). Glycopeptides displaying relatively high molecular masses (large glycans and/or peptides) are less likely to be incorrectly identified due to fewer theoretical glycopeptide candidates (fewer potential false positives) in the higher mass range. In addition, early LC retention time (SO1), high charge (SO3), high glycopeptide mass (SO5), high actual mass error (low mass accuracy, SO6) and high glycan mass (SO8) were search output linked to high *N-*glycoproteome coverage (N4). Teams reporting *N*-glycopeptides with high molecular masses (SO5) more often identified consensus glycopeptides (N5). Finally, low actual mass error (SO6) was, as expected, associated with better identification accuracy.

### High-performance informatics solutions for O-glycoproteomics

Protein Prospector (Team 2) displayed the highest performance in tests scoring the source *O-* glycoproteins (O2) and consensus *O-*glycopeptides (O4) (**Figure 3c**, **Supplementary Table 16**). Conversely, IQ-GPA (Team 1) and glyXtool^MS^ (Team 3) were the best performing software in tests scoring the *O-*glycan compositions (O1) and *O-*glycoproteome coverage (O3), respectively. Overall, Protein Prospector (Team 2, overall score 0.613) and glyXtool^MS^ (Team 3, 0.522) were found to be high-performance software for *O*-glycoproteomics. Amongst the users, four Byonic teams (Team 13, 15, 17, 18, overall score range 0.473-0.701) were ranked in the high-performance band.

Correlation analyses showed that accurate identification of the *O*-glycan compositions (O1) associated with approaches using a focused (narrow) *O-*glycan search space (SS2) and permitting only few missed peptide cleavages (SS7) (**Figure 3d**, **Supplementary Table 18**).

In addition, search strategies permitting incorrect precursor selection (SO4) were commonly used by teams scoring well in the *O*-glycan composition test (O1). Interestingly, employing a broad *O*-glycan search space (SS2) was associated with accurate identification of source *O-* glycoproteins (O2), high *O-*glycoproteome coverage (O3), and better identification of consensus *O-*glycopeptides (O4). Further, teams reporting identifications with low mass error (SO6) scored well in the *O-*glycan composition test (O1), but, notably, at the cost of lower *O-* glycoproteome coverage (O3) and fewer consensus *O-*glycopeptides (O4).

### Search engine-centric analysis

We then explored the impact of different search strategies on the glycoproteomics data output for the popular Byonic search engine used by 11 teams. The Byonic teams employed highly diverse search strategies; except for the common use of decoy/contaminant databases (SS13) and monoisotopic correction (SS14), the search settings varied considerably across these teams (**Figure 4a**). Undoubtedly, this search diversity and different output filtering methods used by the Byonic teams (e.g Byonic score >100, PEP-2D < 0.001, FDR < 1%) contributed to the dramatic variation in reported glycopeptides (**Figure 4b**, **Supplementary Table 1**). Unsurprisingly, therefore, the relative specificity (accuracy) and sensitivity (coverage) scores (established from N1-N6/O1-O5) showed different performance profiles of the Byonic teams particularly for the *O*-glycopeptide analysis (**Figure 4c**, **Supplementary Table 16**). Teams achieving better-than-average sensitivity scores (e.g. Team 15, 17), typically under-performed with respect to specificity. Other teams achieved higher-than-average specificity scores at the cost of sensitivity (e.g. Team 18, 22), confirming the intuitive reciprocal relationship between these performance metrics.

**Figure 4.**
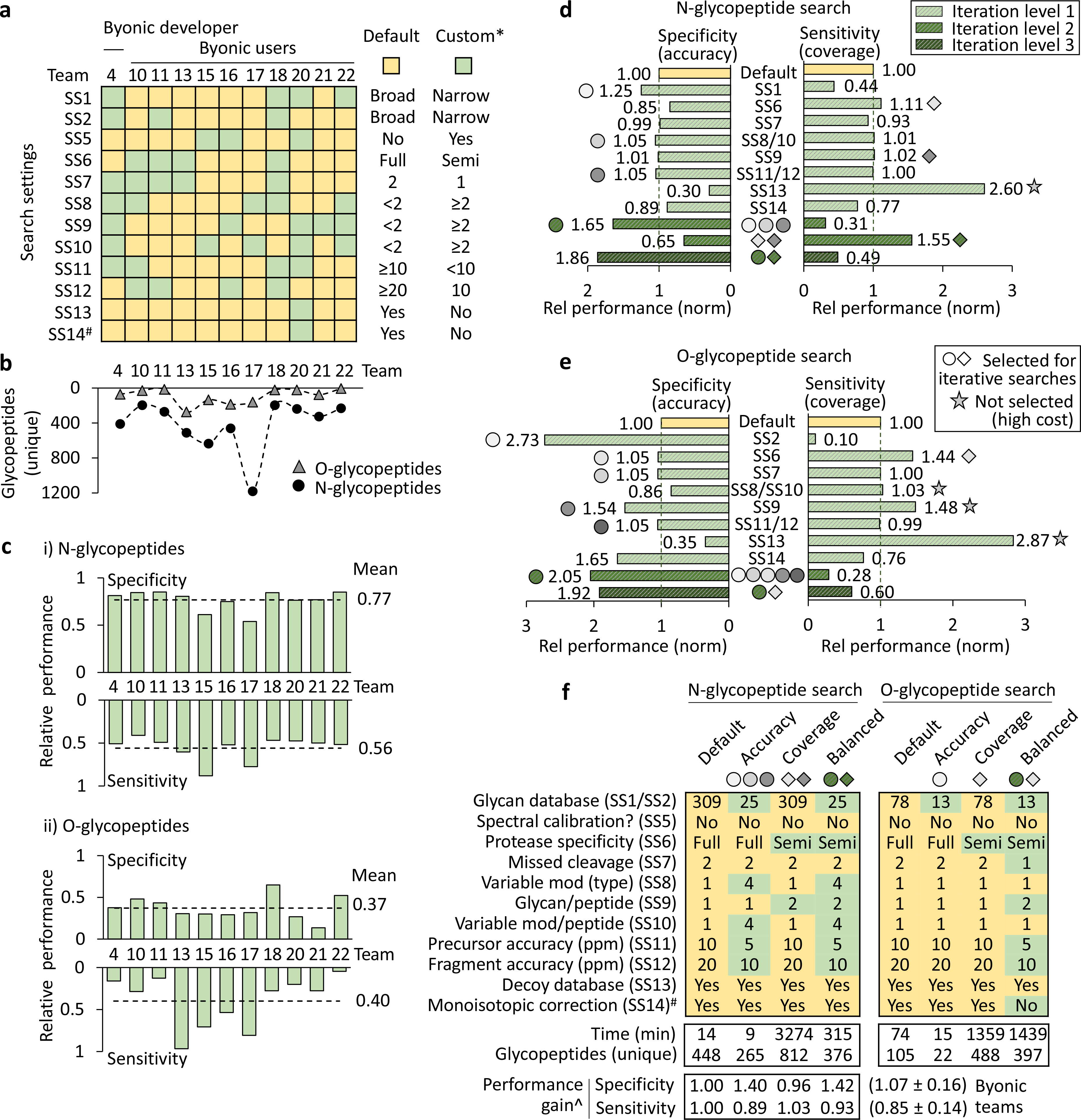
Search engine-centric (Byonic-focused) analysis of search strategies for high-performance glycoproteomics data analysis. **a**. Overview of the search settings employed by Byonic teams. Default: Search strategy used by most teams (yellow). Custom: Variations from the Default search strategy (green). ^#^Data of SS14, a setting not included in the team reports, were adopted from SO4 data. **b**. The glycoproteome coverage (unique glycopeptides, File B) varied amongst Byonic teams. **c**. Specificity (accuracy) and sensitivity (coverage) scores for i) *N*-glycopeptides and ii) *O*-glycopeptides for Byonic teams. **d-e**. Controlled (in-house) searches for *N*- and *O*-glycopeptides using Byonic (File B). Individual search settings were systematically varied (iteration level 1) and output assessed for performance gains (specificity, sensitivity). Search settings showing performance gains (shaded circles/diamonds) without unacceptable costs in specificity (SS13) or search time (SS8/SS10, SS9) (grey stars) were collectively tested for synergy (iteration level 2-3, dark green). **f**. Recommended Byonic-centric search strategies for “high accuracy”, “high coverage” and “balanced” (accuracy><coverage) glycoproteomics data analysis. ^The recommended search strategies showed relative performance gains as determined using an independent glycoprotein-centric score (**Supplementary Table 19b**). Search time and glycoproteome coverage (unique glycopeptides) are also indicated.

The individual search variables were then investigated through a series of controlled (in-house) searches using Byonic. For this purpose, the search settings were systematically varied from the “Default” search strategy used by most teams while keeping other parameters constant. Several search settings showed performance gains in terms of improved specificity (e.g. literature-guided narrow glycan search space, SS1-SS2) or sensitivity (e.g. decoy database disabled, SS13), but often at the expense of other performance characteristics (**Figure 4d-e**, **Supplementary Table 19a**). While reduced sensitivity (glycoproteome coverage) may be an acceptable compromise for higher specificity (identification accuracy), the opposite arguably does not hold true. Thus, the significant sensitivity gains and concomitant loss in specificity achieved by disabling the decoy database (SS13), did not benefit the data analysis. Instead, increasing the permitted glycans per peptide (SS9), tightening the allowed mass error (SS11-SS12) and relaxing the protease specificity benefitted both specificity and sensitivity. Search settings that showed cost-less performance gains were combined for subsequent rounds of iterative searches. Importantly, these efforts led to improved “high accuracy”, “high coverage” and “balanced” (accuracy><coverage) search strategies for *N*- and *O*-glycoproteomics (**Figure 4f**). None of the Byonic teams had utilised these combinations of search settings. When assessed using the independent glycoprotein-centric scoring method, these three recommended search strategies showed improved performance (specificity, sensitivity) relatively to the Default strategy and strategies used by Byonic teams (**Supplementary Table 19b**). Notably, the high coverage searches dramatically expanded the search time, a metric here not considered beyond logistical constraints.

Finally, we explored the performance of different fragmentation methods by systematically varying the spectral input (HCD/EThcD/CID) in Byonic while keeping search settings and output filtering constant. Highest performance was achieved when HCD and EThcD were jointly searched (**Supplementary Table 19c**). Our analysis also suggested that low-resolution CID data do not benefit the Byonic search performance when used alone or with HCD/EThcD data, an observation supported by the Byonic team comparison (**Supplementary Table 19d**).

## Discussion

This is the first community-driven study aiming to objectively discern the performance of current informatics solutions for glycoproteomics data analysis. Excitingly, several high-performance glycoproteomics software and search strategies were identified. Amongst the 9 developer teams, Protein Prospector (Team 2) was identified as the top performing software for both *N*- and *O*-glycoproteomics. Byonic (Team 4) also displayed high-performance for *N*-glycopeptide data analysis, and while this developer only demonstrated moderate performance for *O*-glycoproteomics, four Byonic user teams (Team 13, 15, 17, 18) displayed the highest performance for *O*-glycopeptide data analysis. Protein Prospector^51^ and Byonic^33^, developed 10-20 years ago, have pioneered the glycopeptide informatics field and are search engines already commonly used in glycoproteomics^8, 31, 33^.

Protein Prospector is an academic (free) tool recognised for its ability to identify modified peptides and modification site(s) from LC-MS/MS data using a probability-difference based scoring system^31^. Protein Prospector is often a preferred search engine in studies addressing the challenging site annotation of *O*-glycopeptides in particular when ET(hc)D-MS/MS data are available^5^. However, Protein Prospector does not estimate the FDR of the glycan components of glycopeptides, which it regards as non-descript PTMs with an exact mass, and the software may appear less user-friendly than competing tools.

Facilitated by a user-friendly interface, precise spectral annotation and useful output reports of identified glycopeptides/proteins, the commercial Byonic search engine has gained considerable popularity (as illustrated herein) as it enables relatively straightforward identification of peptides with known and unknown modifications including glycosylation from different MS/MS data. Byonic features useful fine control options that enable tailored glycopeptide searches and post-search filtering of output based on prior knowledge. Byonic scores and annotates multiple types of glycopeptide fragments to deduce the peptide carrier, glycan and modification site, but the FDRs, calculated identically for non-glycosylated peptides and glycopeptides, primarily addresses the correctness of the peptide rather than the glycan and the site localisation^33^.

Notably, GlycoPAT^37^ (Team 8) and glyXtool^MS^ ^32^ (Team 3) were in our study also identified as high-performance *N*- and *O*-glycoproteomics software, respectively. Furthermore, IQ-GPA^30^ (Team 1) demonstrated merit for both *N*- and *O*-glycopeptide data analysis. While all three software handle high-resolution HCD-, ETciD-, and EThcD-MS/MS data, IQ-GPA and GlycoPAT also identify glycopeptides based on high-resolution CID-MS/MS data and apply post-search filtering based on advanced peptide and glycan decoy methods to estimate both peptide and glycan FDRs of glycopeptide candidates. The software glyXtool^MS^ instead uses oxonium ions, Y_1_-ions (peptide-HexNAc), other glycopeptide-specific fragments, and peptide-specific b-/y-ions to control FDR. Excitingly, these three academic tools were recently developed (< 5 years ago), and thus, hold a considerable potential in the field.

We used both team-wide and search engine-centric approaches to uncover performance-associated search variables for glycoproteomics data analysis. The team-wide correlation analyses revealed many search settings and search output linked to performance. Backed by robust statistics, these “universal” relationships existing across search engines will widely benefit glycoproteomics software developers and users aiming to improve *N*- and *O*-glycopeptide data analysis (**Table 2**). This knowledge may aid tackling existing challenges in glycoproteomics, amongst the most critical, reducing the FDR of glycopeptide candidates carrying glycans with similar (e.g. NeuAc-R *versus* Fuc_2_-R, Δm = 1.0204 Da) or identical (e.g. NeuAc_1_Hex_1_-R *versus* NeuGc_1_Fuc_1_-R, Δm = 0 Da) masses. Our study indeed confirms that glycopeptides displaying such “difficult-to-identify” features (NeuAc, NeuGc, multi-Fuc, Met oxidation, Cys carbamidomethylation) are frequently mis-annotated with current search engines (**Extended Data Figure 5**). We therefore recommend that efforts should be invested in improving tools to allow for accurate identification of such challenging glycopeptides.

**Table 2.**
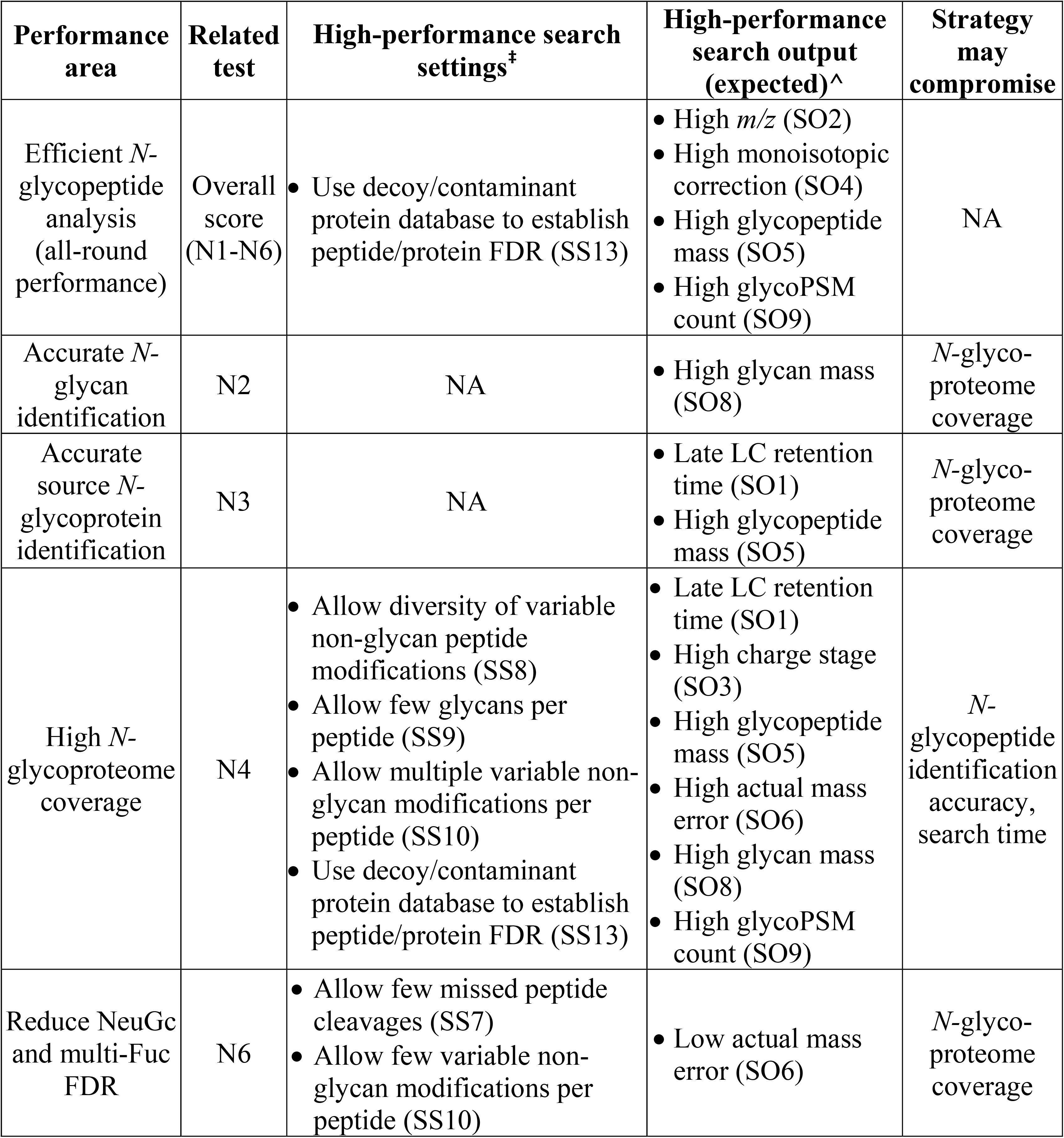
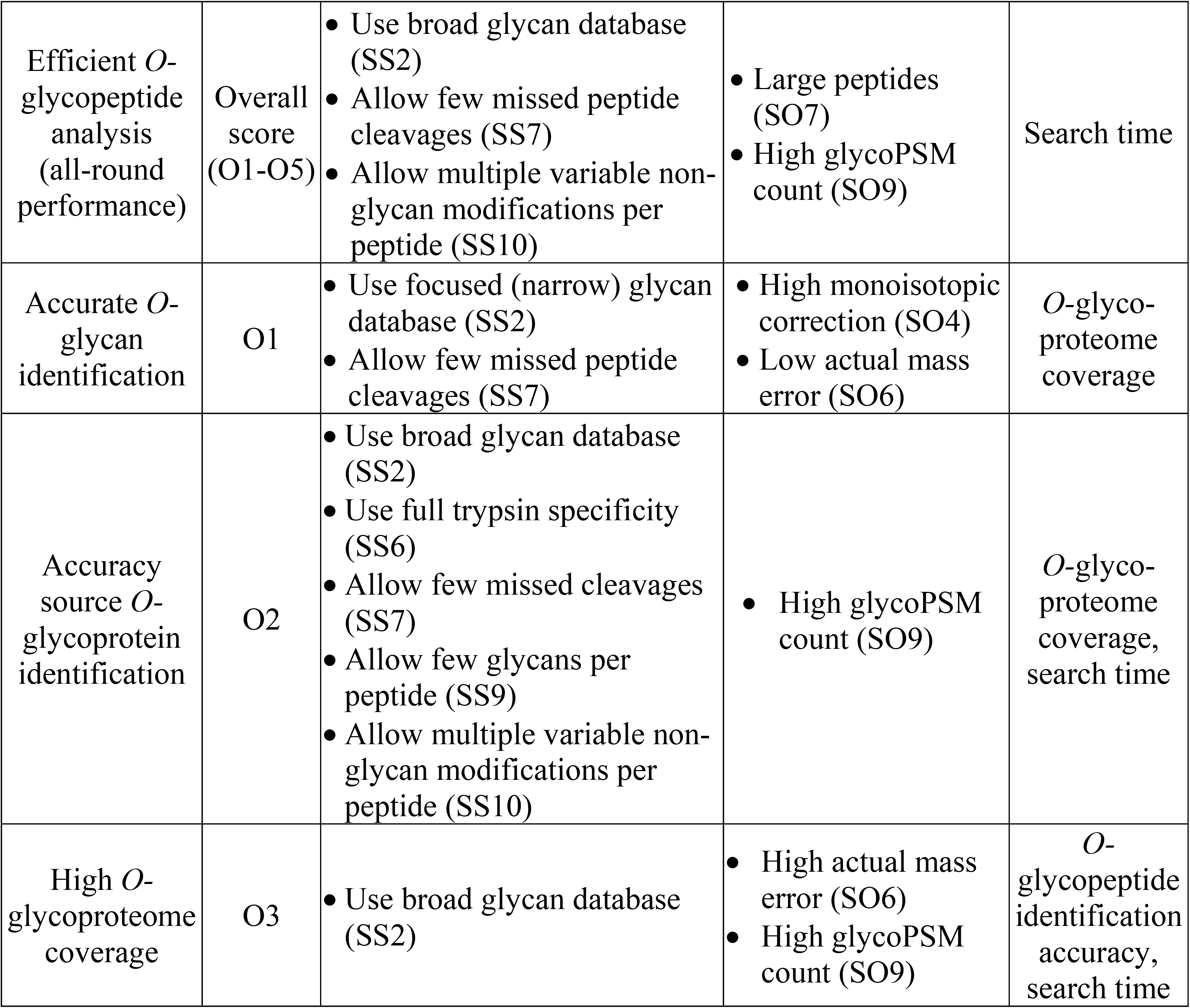
Overview of software-independent search variables important for high-performance glycoproteomics data analysis (see **Figure 3b,d**, Table 1, Supplementary Table 18 for study variables and associations). Only search variables closely associated with high performance (≥3 statistical tests) have been included. ^‡^Software-independent search settings that may guide improved glycoproteomics search strategies. ^Search output expected from high-performance glycoproteomics data analysis. This information may also aid post-search filtering of glycopeptide data. The possible compromise of selected search strategies on the overall glycoproteomics performance is indicated. NA, not applicable.

| Performance area | Related test | High-performance search settings<sup>‡</sup> | High-performance search output (expected)<sup>^</sup> | Strategy may compromise |
| --- | --- | --- | --- | --- |
| Efficient *N*-glycopeptide analysis (all-round performance) | Overall score (N1-N6) | • Use decoy/contaminant protein database to establish peptide/protein FDR (SS13) | • High *m/z* (SO2)<br>• High monoisotopic correction (SO4)<br>• High glycopeptide mass (SO5)<br>• High glycoPSM count (SO9) | NA |
| Accurate *N*-glycan identification | N2 | NA | • High glycan mass (SO8) | *N*-glycoproteome coverage |
| Accurate source *N*-glycoprotein identification | N3 | NA | • Late LC retention time (SO1)<br>• High glycopeptide mass (SO5) | *N*-glycoproteome coverage |
| High *N*-glycoproteome coverage | N4 | • Allow diversity of variable non-glycan peptide modifications (SS8)<br>• Allow few glycans per peptide (SS9)<br>• Allow multiple variable non-glycan modifications per peptide (SS10)<br>• Use decoy/contaminant protein database to establish peptide/protein FDR (SS13) | • Late LC retention time (SO1)<br>• High charge stage (SO3)<br>• High glycopeptide mass (SO5)<br>• High actual mass error (SO6)<br>• High glycan mass (SO8)<br>• High glycoPSM count (SO9) | *N*-glycopeptide identification accuracy, search time |
| Reduce NeuGc and multi-Fuc FDR | N6 | • Allow few missed peptide cleavages (SS7)<br>• Allow few variable non-glycan modifications per peptide (SS10) | • Low actual mass error (SO6) | *N*-glycoproteome coverage |
| Efficient *O*-glycopeptide analysis (all-round performance) | Overall score (O1-O5) | • Use broad glycan database (SS2)<br>• Allow few missed peptide cleavages (SS7)<br>• Allow multiple variable non-glycan modifications per peptide (SS10) | • Large peptides (SO7)<br>• High glycoPSM count (SO9) | Search time |
| Accurate *O*-glycan identification | O1 | • Use focused (narrow) glycan database (SS2)<br>• Allow few missed peptide cleavages (SS7) | • High monoisotopic correction (SO4)<br>• Low actual mass error (SO6) | *O*-glycoproteome coverage |
| Accuracy source *O*-glycoprotein identification | O2 | • Use broad glycan database (SS2)<br>• Use full trypsin specificity (SS6)<br>• Allow few missed cleavages (SS7)<br>• Allow few glycans per peptide (SS9)<br>• Allow multiple variable non-glycan modifications per peptide (SS10) | • High glycoPSM count (SO9) | *O*-glycoproteome coverage, search time |
| High *O*-glycoproteome coverage | O3 | • Use broad glycan database (SS2) | • High actual mass error (SO6)<br>• High glycoPSM count (SO9) | *O*-glycopeptide identification accuracy, search time |

Meanwhile, our search engine-centric approach involving systematic Byonic searches revealed search settings impacting the performance of this widely used search engine and highlighted that specificity (accuracy) and sensitivity (coverage) are competing performance characteristics challenging to achieve in a single search. This suggests that glycoproteomics data may benefit from being interrogated using multiple orthogonal search strategies that are subsequently combined or by approaches that strike a balance between accuracy and coverage. To this end, we here recommend a set of improved “high accuracy”, “high coverage” and “balanced” search strategies that should be selected (and further tailored/optimised) according to the sample and research question being investigated (**Figure 4f**).

Although not a focus here, our study also showed that the search strategy dramatically impacts the search time. While the spectral input type and data output filtering represent other critical variables that also need further exploration, our study indicates that HCD- and EThcD are currently the most informative spectral types in glycoproteomics, and that knowledge-guided filtering and curation of data output is critically required to lower FDRs.

The study also highlighted the significant informatics challenges still associated with large-scale glycopeptide data analysis as illustrated by the discrepant reporting of glycopeptides across teams. Notably, high discordance of reported glycopeptides was even found between participants using the same software, confirming that search variables other than the search engine also substantially impact the glycopeptide data analysis. While the 10 Byonic user teams reported a marginally higher rate of consensus glycopeptides, their spread in terms of search output data, reported glycoPSMs and overall performance scores was of similar magnitude as the variance observed amongst other user teams. Whilst the (self-reported) team experience in glycoproteomics was not found to be an accurate predictor of team scoring and ranking, the variability in the spectral data input, search settings and, importantly, at the post-search filtering stage were identified as key factors contributing to the discrepant reports. Concertedly, these observations point to the importance of using both the most informative spectral data, powerful search engines, tailored search settings and knowledge-driven post-search filtering to achieve high-performance glycoproteomics data analysis.

Despite the considerable team-to-team variation, this study produced consensus lists of 163 *N-*glycopeptides and 23 *O*-glycopeptides from serum glycoproteins commonly reported by teams. Importantly, these high-confidence glycopeptides carried biosynthetically-related glycans that were devoid of NeuGc and poor in multi-Fuc features in line with literature^23, 40^ and mapped to known high-abundance serum proteins^4, 23, 39, 41, 42^. The consensus lists have been made publicly available (GlyConnect ID 2943) as they form an important reference for future studies of the human serum glycoproteome.

The study design including the sample type/preparation and data collection method was chosen to mimic conditions typically encountered in glycoproteomics while also aiming to accommodate most informatic solutions and appeal to users in the field. Multiple orthogonal performance tests and separate validation were applied to ensure a fair and holistic scoring of search engines and teams. Despite these efforts, it cannot be ruled out that some software or users may have been unintentionally disadvantaged and/or excluded by the chosen experimental design and scoring system. The team scoring and ranking should be viewed in light of these constraints and limitations common to most community-based comparison studies founded on communal data.

In addition to reporting on the peptide and glycan components of identified glycopeptides, teams were requested to report on site(s) of modification where possible. Since most tryptic *N*-glycopeptides only comprise a single sequon, site localisation is primarily a challenge related to *O*-glycoproteomics^5, 16^. Most teams indeed returned data of the *O*-glycosylation site(s), but due to highly discrepant and often inconclusive reporting of sites and a paucity of literature on serum *O*-glycosylation sites, we were unable to score glycosylation site localisation.

Most software currently available for glycoproteomics data analysis participated in this study. However, several glycopeptide search engines e.g. pGlyco^52^, MSFragger-Glyco^53^, O-Pair Search^54^, and StrucGP^55^ were unfortunately not represented due to LC-MS/MS data incompatibility or due to their development after the study period. Thus, this study is essentially a snapshot of the performance of software available at the time the data analysis was performed. Highlighting the rapid progress in glycoproteome informatics, most of the software solutions participating in this study have been improved and new versions released after the evaluation period. For example, GPQuest v2.0, GlycoPAT v1.0 and Protein Prospector v5.20.23 tested herein have been superseded by more recent versions i.e. GPQuest v2.1, GlycoPAT v2.0 and Protein Prospector v.6.2.2. Thus, a limitation of this study is that newer tools are available at the time of publication that were not compared in our analysis. Follow-up studies comparing the performance of these latest glycoproteomics software upgrades and informatics solutions not included in this study are therefore warranted. Beyond testing the ability of participants to identify the peptide and glycan components of glycopeptides from glycoproteomics data, such future comparative studies should ideally also test the ability to accurately quantify (relative, absolute) and report on modification sites of identified glycopeptides and could explore other relevant parameters not addressed herein including the use of alternative proteases, TMT-labelling, and stepped-HCD-MS/MS data amongst other experimental conditions gaining popularity in glycoproteomics.

In summary, this community study has documented that the field has several high-performance informatics solutions available for glycoproteomics data analysis and has elucidated key performance-associated search strategies that will serve to guide developers and users of glycoproteomics software.

## Supporting information

Supplementary Information containing Extended Methods

## Acknowledgements

Drs Rosa Viner and Sergei Snovida (Thermo Fisher Scientific) are thanked for providing high-quality LC-MS/MS data. Dr Krishnatej Nishtala is thanked for aiding the data analysis. Drs Catherine Hayes, Julien Mariethoz and Frederique Lisacek are thanked for informatics assistance. RK was supported by an Early Career Fellowship (Cancer Institute NSW ECF181259). DK was supported by an Australian Research Council Future Fellowship (FT160100344). GP was funded by FAPESP (n° 2018/15549-1). MTA was supported by a Macquarie University Safety Net Grant. DK and NHP were supported by the Australian Research Council Centre of Excellence in Nanoscale Biophotonics (CE140100003).

## Author Contributions Statement

Conception: NHP, MTA. Design: RK, DK, KK, GL, KFM, GP, JZ, JSY, SMH, NHP, MTA. Data acquisition: All participants (Team 1-22). Data analysis: RK, AC, BLP, GS, MTA. Data interpretation: RK, AC, MTA. Creation of new software used in the work: All developers (Team 1-9). Manuscript writing/editing: RK, AC, DK, KK, GL, KFM, GP, JZ, JSY, SMH, NHP, MTA. All authors have approved the manuscript.

## Online Methods

Key methodological details are provided below. Please consult the **Extended methods** for an exhaustive description of methods employed.

### Study design and participants

Nine developer and 13 user teams completed the study (**Supplementary Table 1**). Teams received two glycoproteomics data files (File A-B) and reported identified *N*- and *O*-glycopeptides using a common reporting template (PXD024101, PRIDE). User teams were free to use any search engine(s) at their disposal including manual annotation/filtering of search output. Developers returned the identified glycopeptides directly from their own software without manual post-search filtering. The relative team performance was therefore compared within (not between) the developer and user groups.

### Study sample

A synthetic *N*-glycopeptide (52 fmol) from human vitamin K-dependent protein C (EVFVH<u>P</u>NYSK, Hex_5_HexNAc_4_NeuAc_2_) was spiked into human serum (product #31876, Thermo Fisher Scientific), the mixture digested using trypsin, and glycopeptides enriched.

### Mass spectrometry

The enriched glycopeptide mixture was analysed twice using two similar LC-MS/MS acquisition methods generating File A-B. For both files, peptides were separated using C_18_ reversed-phase nanoLC and detected on a Thermo Scientific™ Orbitrap Fusion™ Lumos™ Tribrid™ mass spectrometer using high-resolution MS1 and data-dependent HCD-MS/MS acquisition. Product-dependent (oxonium ion) triggered ETciD-/CID-MS/MS (Orbitrap) and EThcD-/CID-MS/MS (Orbitrap/ion trap) events of glycopeptide precursors were scheduled for File A-B, respectively. Participants received File A-B as .raw/.mgf files.

### Search instructions and reporting template

Teams used a fixed protein search space (human proteome, 20,231 UniProtKB reviewed sequences), but could freely choose the glycan search space. Teams were instructed neither to include xylose nor any glycan substitutions in the glycan search space. The participants reported their team details, identification strategy and identified glycopeptides in a common reporting template. All reports were carefully checked for compliance to the study guidelines to enable fair comparisons between teams.

### Compilation and comparison of team data

Data from the returned reports were compiled (**Supplementary Table 1-2**) including the reported glycopeptides (**Supplementary Table 3**). Unique identifiers (IDs) for the reported glycopeptides and their glycan compositions and source glycoproteins were generated to enable team-to-team comparisons using pivot tables in Excel.

### Glycoprofiling of select serum N-glycoproteins

The site-specific *N*-glycosylation of select glycoproteins including human alpha-1-antitrypsin (A1AT, P01009), ceruloplasmin (CP, P00450), haptoglobin (HP, P00738) and immunoglobulin G1 (IgG1, P01857) was profiled using AUC-based MS1 quantitation.

### Search engine-centric analysis and optimisation of Byonic search strategies

Deep search engine-centric analyses were performed through a series of controlled in-house searches using Byonic v3.9.4 (Protein Metrics Inc., CA, USA). Search settings or spectral data input were systematically varied and the output scored. Search settings leading to relative performance gains were iteratively used for additional searches.

### Performance testing of teams and software

Relative performance for glycopeptide data analysis was determined via three different methods. In short, teams were comprehensively scored and ranked based on overall performance scores established from multiple complementary performance tests (N1-N6, O1-O5). The performance tests included the synthetic *N-*glycopeptide test (N1), the glycan composition test (N2, O1), the source glycoprotein test (N3, O2), the glycoproteome coverage test (N4, O3), the commonly reported (consensus) glycopeptide test (N5, O4), and the NeuGc/multi-Fuc glycopeptide test (N6, O5). The team scoring was validated using an independent glycoprotein-centric assessment method that scored the quantitative match of glycoPSMs reported by teams to the actual site-specific *N*-glycosylation of selected glycoproteins (A1AT, CP, HP, IgG1). For the search engine-centric analysis of the Byonic search strategies, performance was evaluated based on relative specificity and sensitive scores derived from the performance tests (N1-N6, O1-O5).

### Statistical analysis

Scores from each performance test (N1-N6, O1-O5) and the overall team performance scores were tested for associations with the search settings (SS1-SS13) and search output (SO1-SO9) using seven statistical methods^56^. Only strong associations observed across at least three different statistical methods were considered. Further, unpaired two-sided t-tests were applied as indicated. The confidence interval was 95% and statistical significance indicated as \**p* < 0.05, \*\**p* < 0.01 and \*\*\**p* < 0.001. Pearson correlations were used to quantitatively compare the observed site-specific *N*-glycan distribution of four glycoproteins in the sample to the literature and to validate the team scoring via the glycoprotein-centric scores.

## Data Availability Statement

This study contains extended data figures and other supplementary information. Multiple figures/tables have associated raw data (**Figure 1-4**, **Table 1-2**, **Extended Data Figure 1-4**, **8-10**). The supporting information includes: 1) **Extended Data Figure 1-10,** 2) Supplementary information containing **Extended methods** (pdf) and 3) **Supplementary Table 1–19** (Microsoft Excel). Further, the LC-MS/MS raw data (File A-B), reporting template, and deidentified but otherwise unredacted team reports are available via ProteomeXchange (PXD024101). Username, Password: YLk2wW1P. The consensus glycopeptides are available via the GlyConnect resource of the Glycomics@ExPASy collection hosted at SIB - Swiss Institute of Bioinformatics (GlyConnect Reference ID 2943).

## Code Availability Statement

Developers (Team 1-9) used their own software to complete this study. All participants including the developers detailed how software were handled and how data were generated. Additionally, developers detailed in their reports how their software could be tested and data validated by the study committee. While the codes for the developer software (commercial/academic origins) have not been released as part of this work, all team reports underpinning this study have been released (see Data Availability Statement).

---

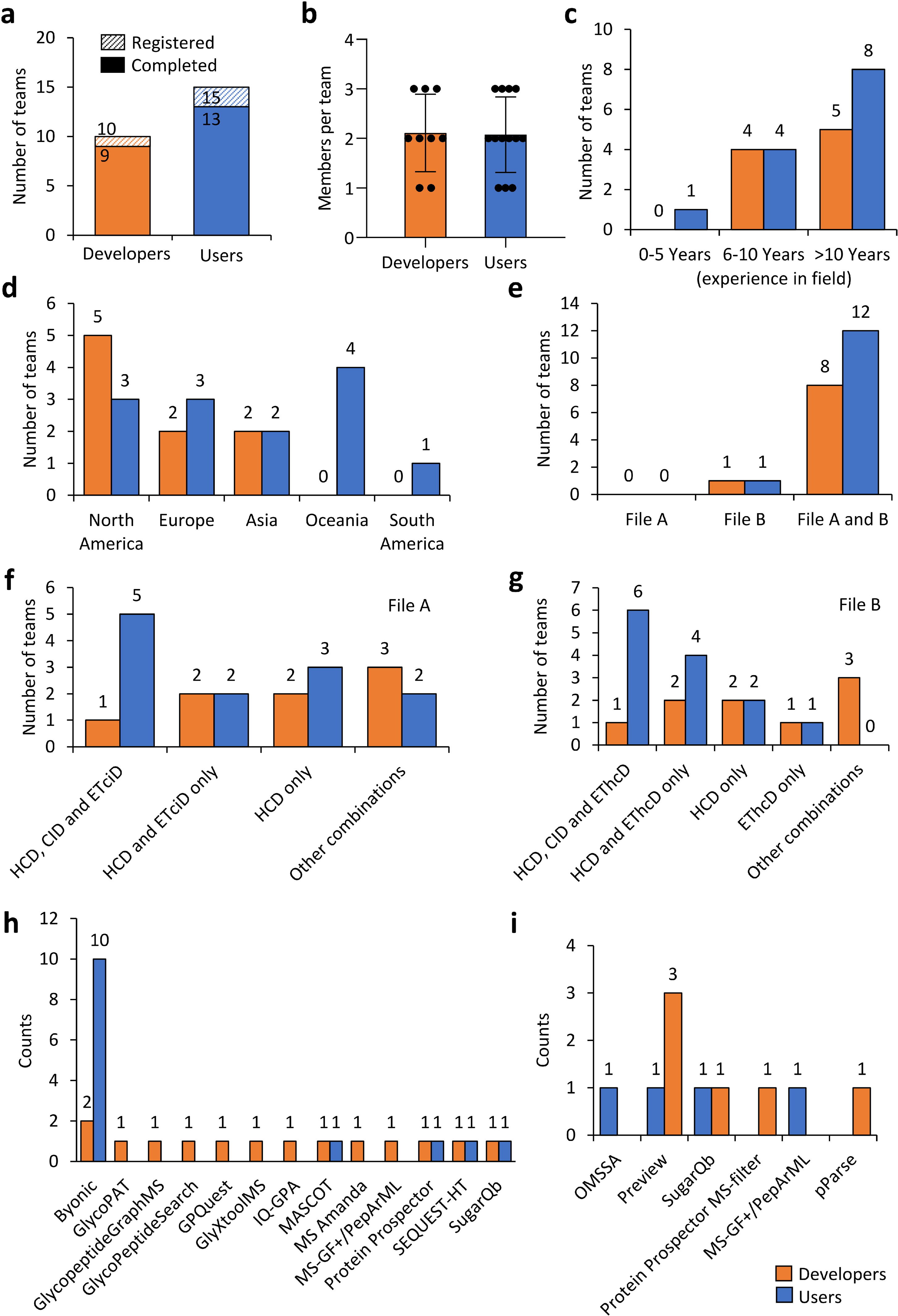

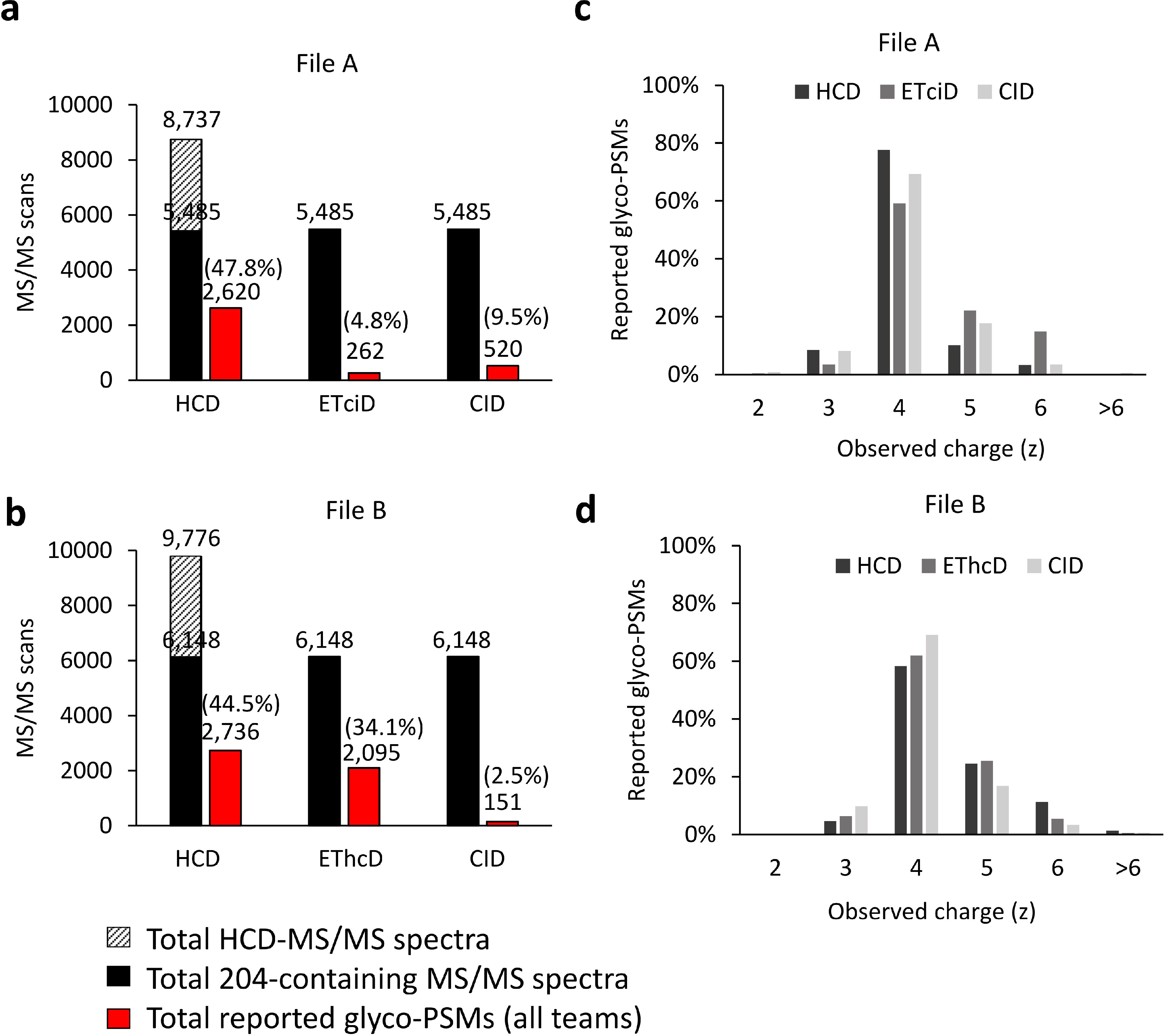

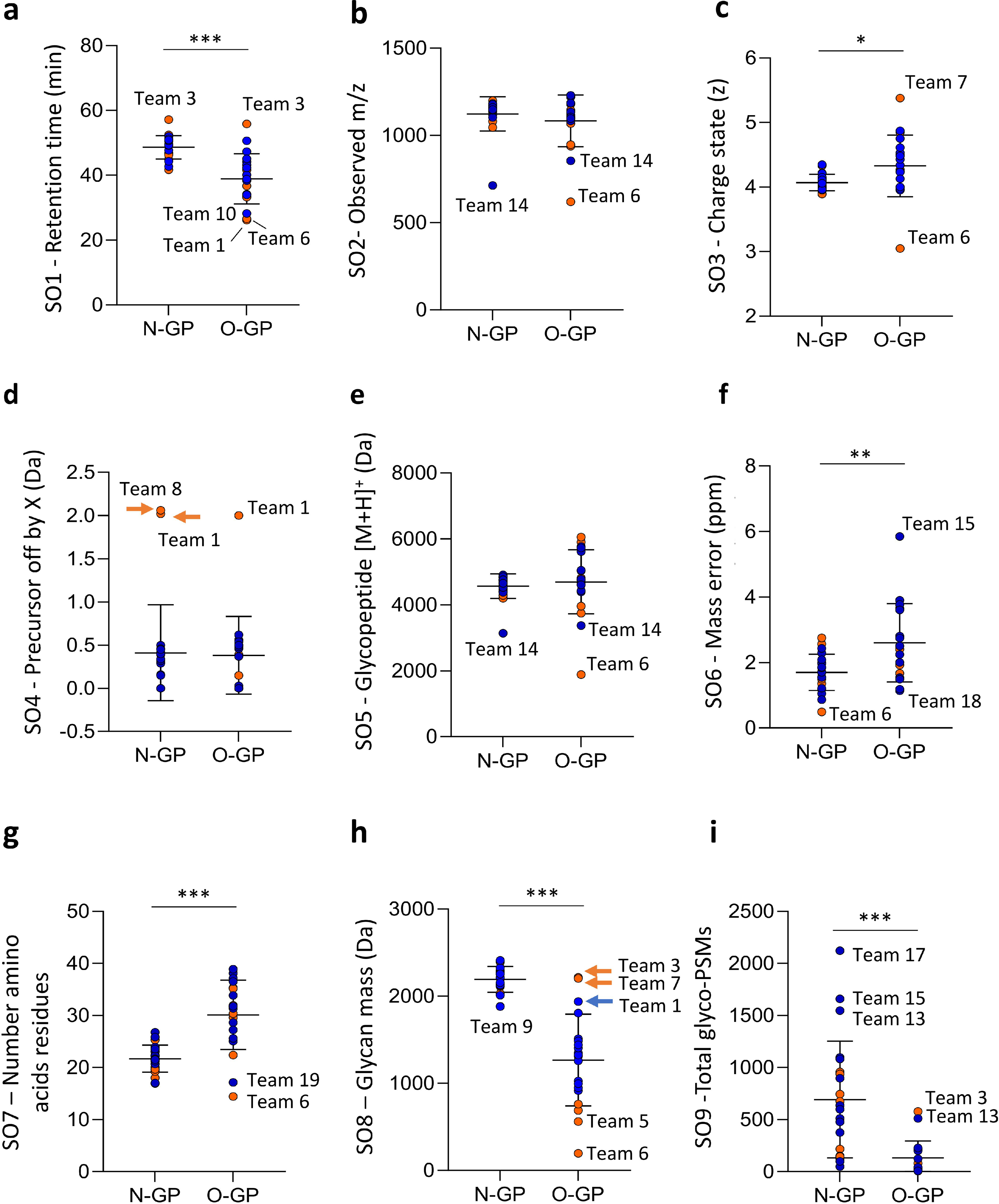

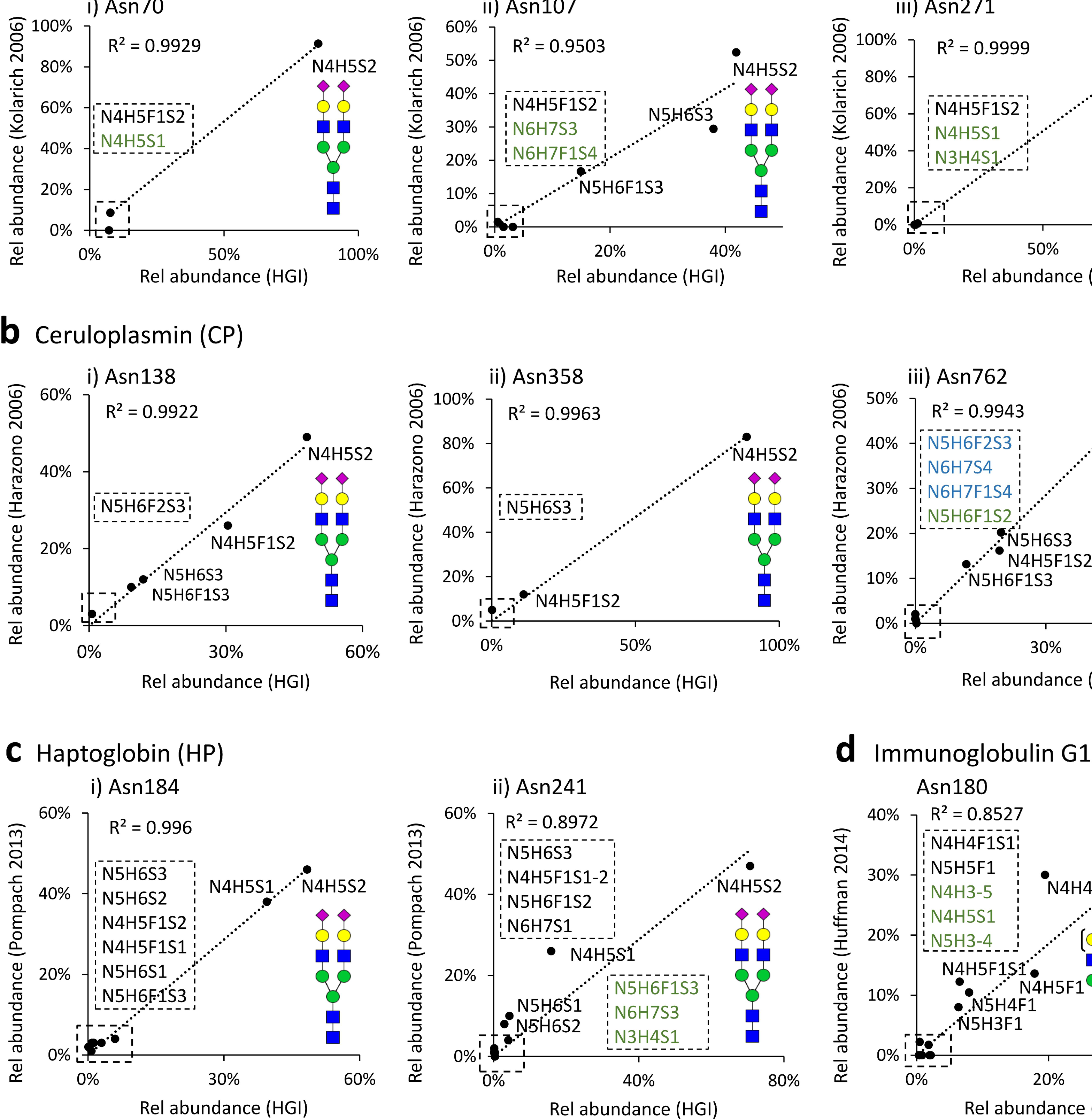

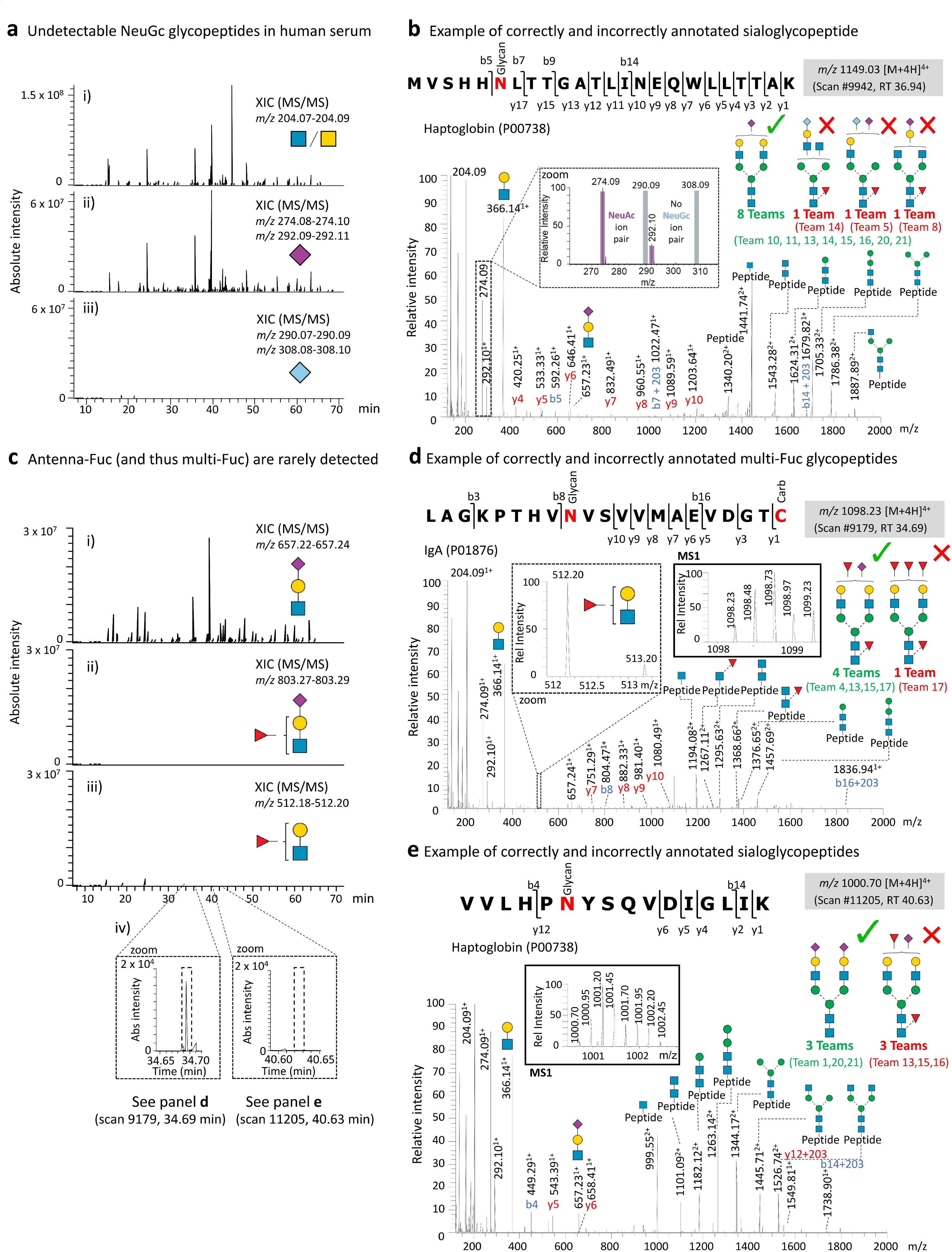

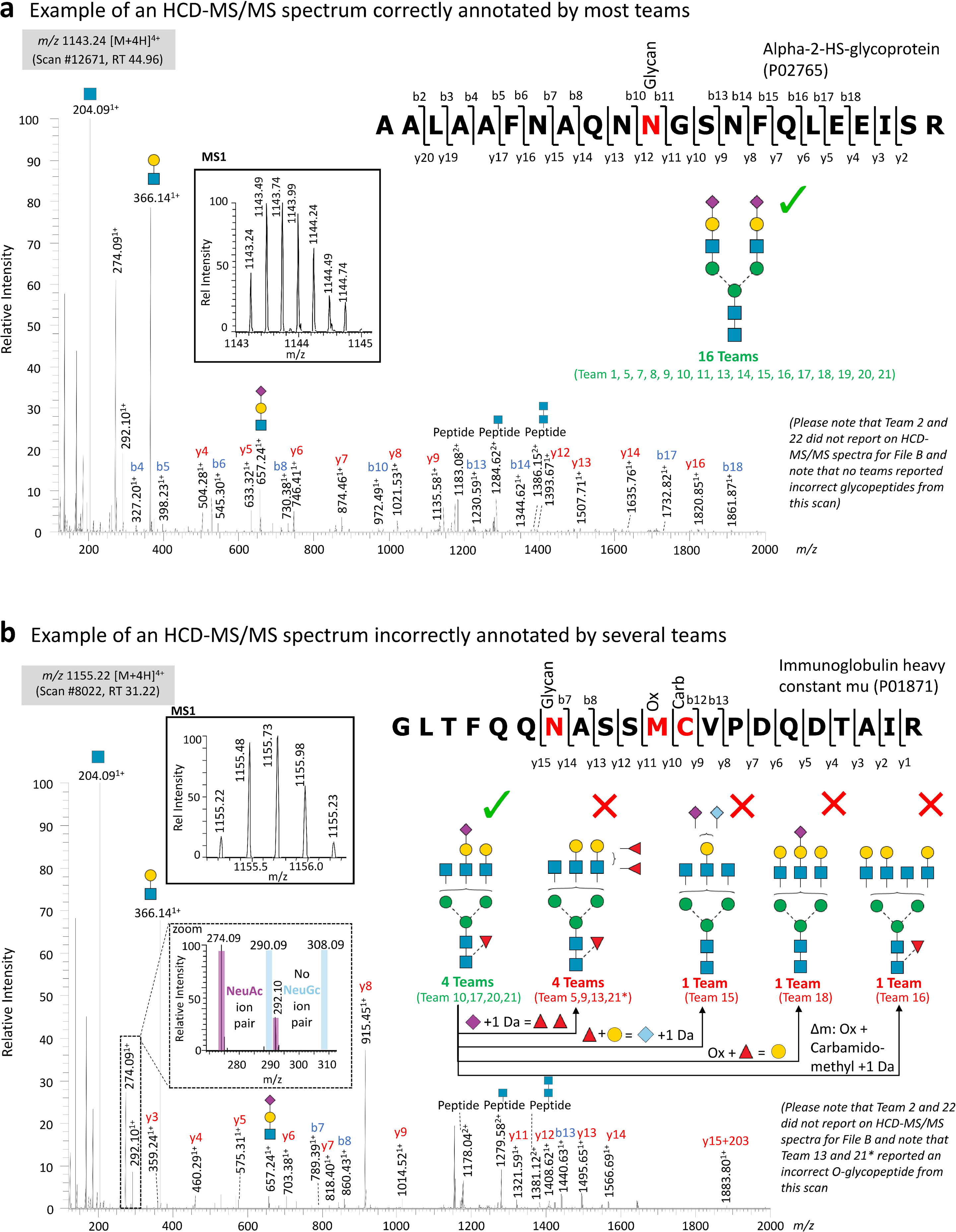

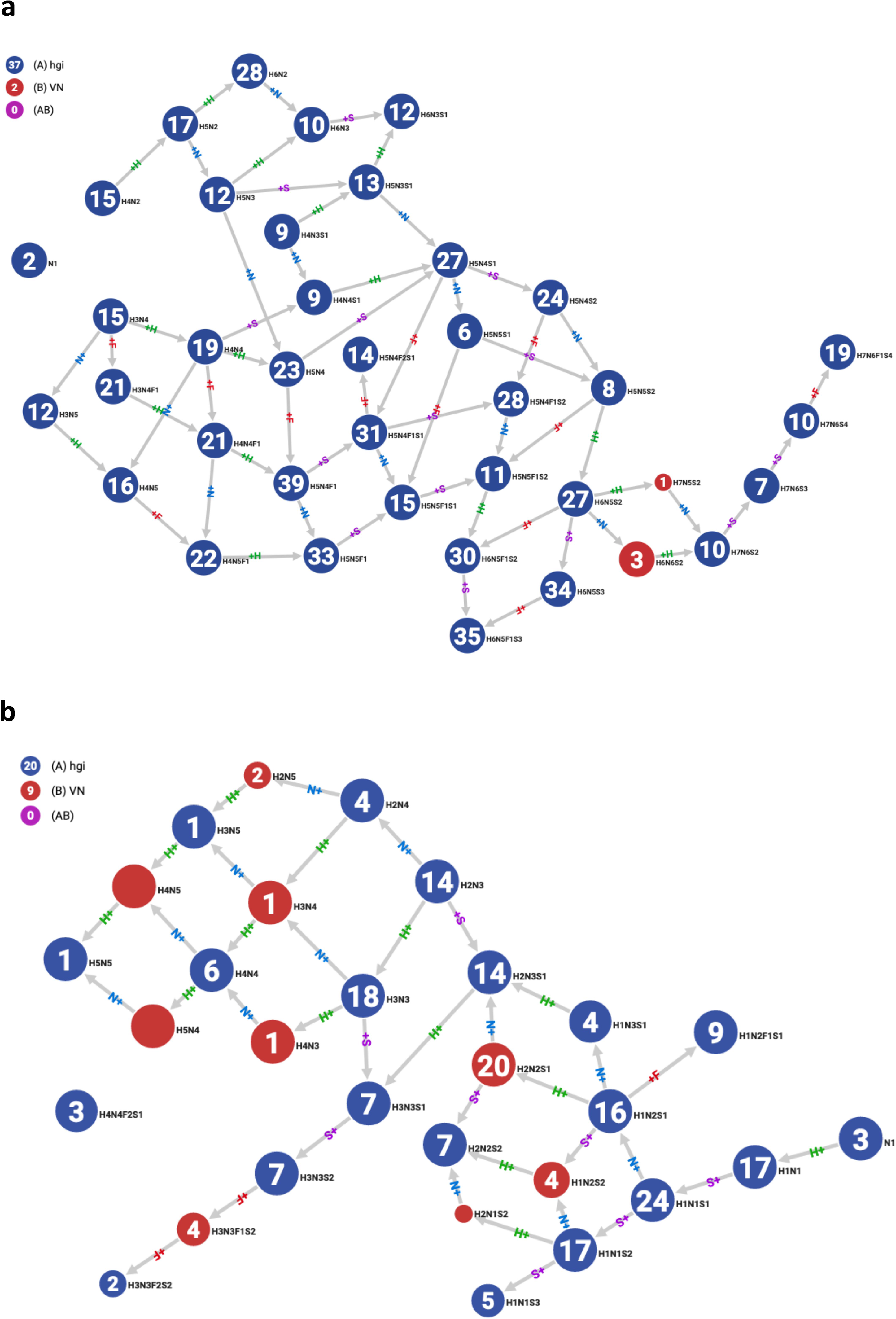

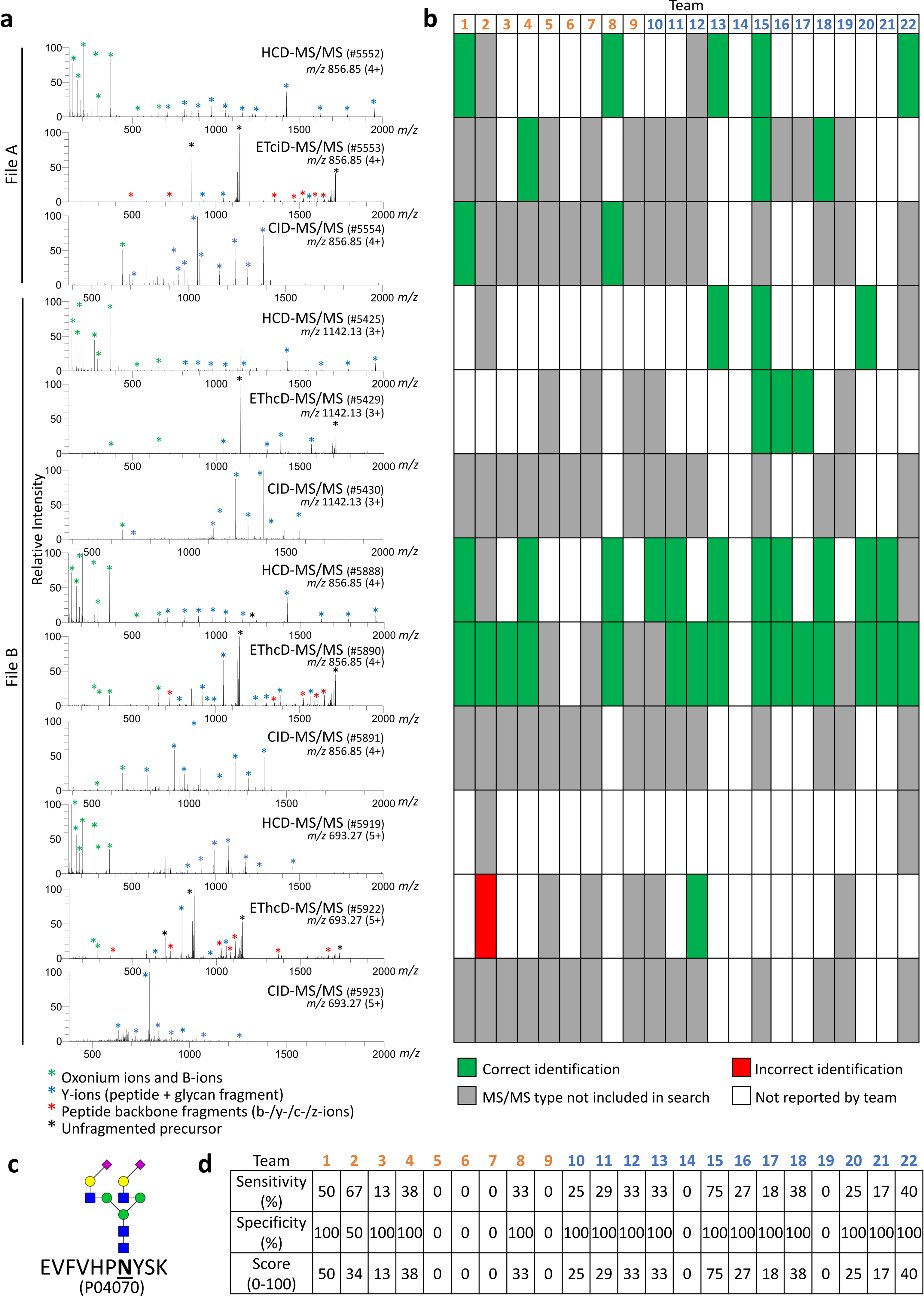

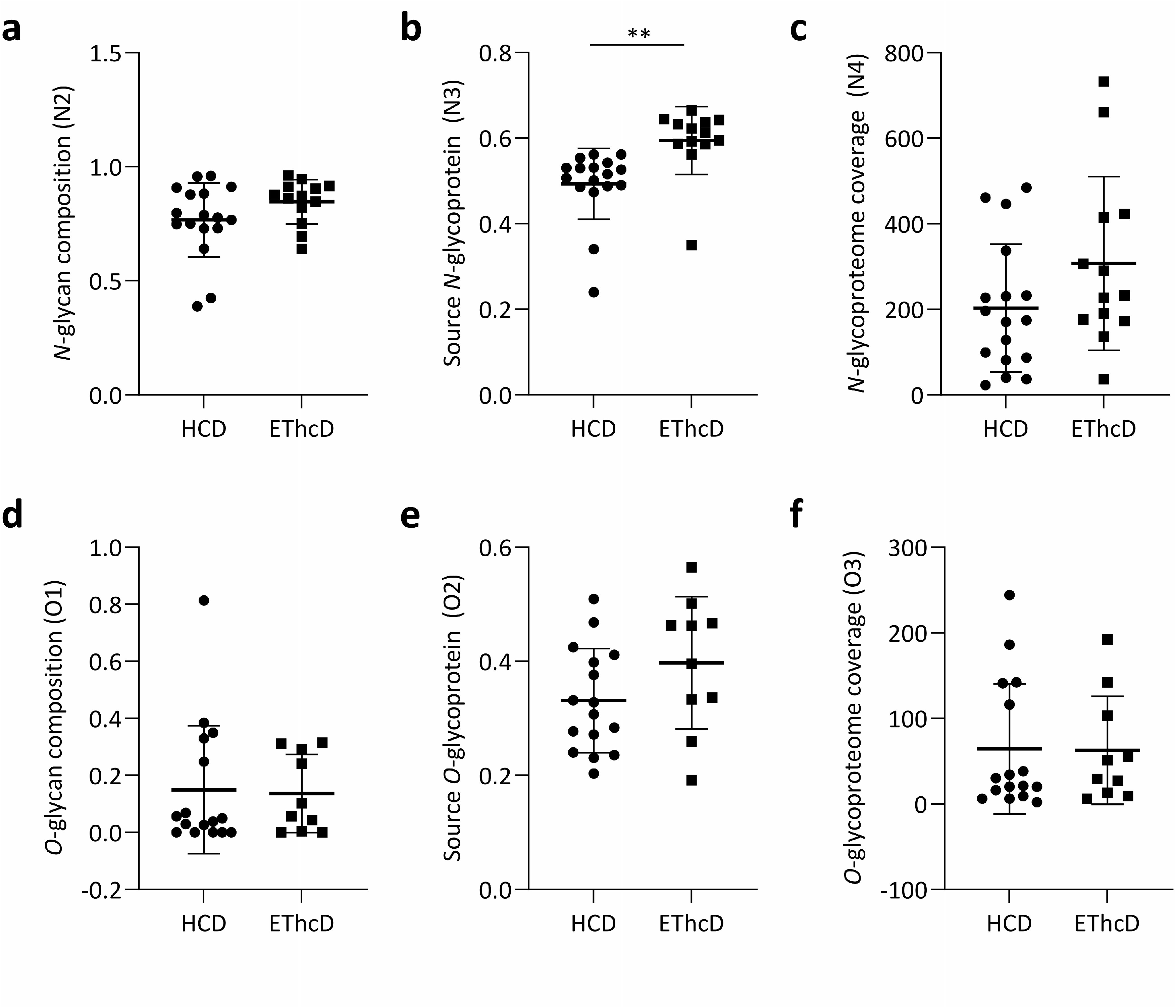

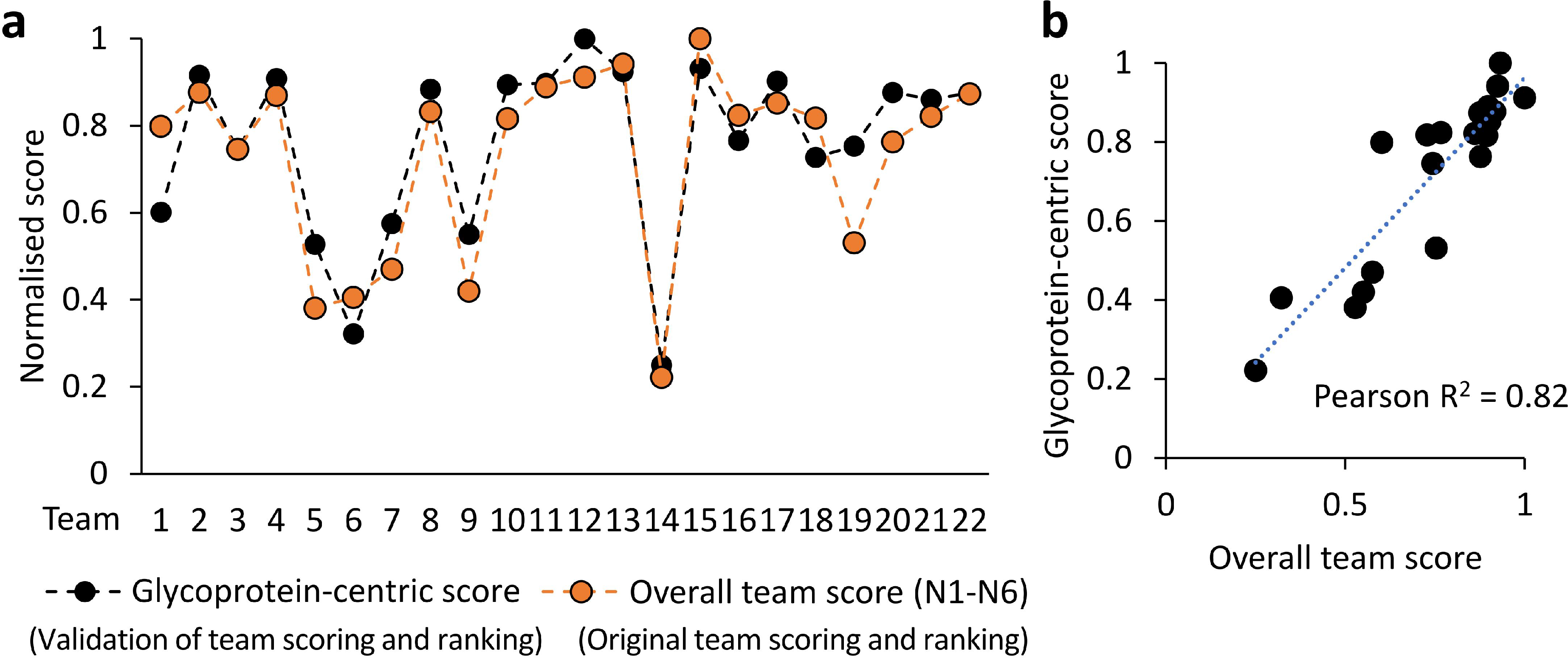

