## Supplementary Information containing Extended Methods for "Community Evaluation of Glycoproteomics Informatics Solutions Reveals High-Performance Search Strategies of Serum *N*- and *O*-Glycopeptide Data"

for

**Running title**: High-performance search strategies of serum glycoproteomics data

**Keywords**: Glycoproteomics, glycopeptide, *N*-glycosylation, *O*-glycosylation, informatics, software, search engine, mass spectrometry

***Corresponding author**:

Dr Morten Thaysen-Andersen, PhD

Department of Molecular Sciences

Biomolecular Discovery Research Centre

Faculty of Science & Engineering

Macquarie University - Sydney

NSW-2109, Australia

### Extended Methods

*Study design and participants*

Calls to join this study as a developer (academic/commercial) or user of glycoproteomics software were made widely across the proteomics and glycomics community. In total, 25 teams signed up for the study, out of which 22 teams comprising 9 developer and 13 user teams completed the study. All teams identified *N*- and *O*-glycopeptides from two communal glycoproteomics LC-MS/MS data files (File A-B), and reported their findings using a common reporting template (PXD024101, PRIDE repository). While the user teams were guaranteed anonymity, the developers were informed that their software (hence, potentially their identity) would be disclosed upon publication. The user teams were free to use any search engine(s) at their disposal including manual annotation/filtering of search output. Developers returned the identified glycopeptides directly from their own software without manual post-search filtering. The developers employed the following search engines: Team 1: IQ-GPA v2.5[1], Team 2: Protein Prospector v5.20.23[2], Team 3: glyXtool$^{MS}$ v0.1.4[3], Team 4: Byonic v2.16.16[4], Team 5: Sugar Qb[5], Team 6: Glycopeptide Search v2.0alpha[6], Team 7: GlycopeptideGraphMS v1.0/Byonic[7], Team 8: GlycoPAT v2.0[8] and Team 9: GPQuest v2.0[9] (see **Supplementary Table 1**). The relative team performance was compared within (not between) the developer and user groups since these two groups were given slightly different instructions (see above).

*Synthetic N-glycopeptide*

An Asn-building block carrying a disialylated, biantennary *N*-glycan ($Hex_5HexNAc_4NeuAc_2$) was purified from chicken egg yolk powder. Previous studies have confirmed that a disialylated, biantennary *N*-glycan carrying only α2,6-linked NeuAc residues is the major component of the chicken egg yolk hexapeptide[10, 11]. In short, this glycosylated hexapeptide was subjected to extensive proteolysis to generate a glycosylated Asn, which was then converted into a fluorenylmethoxycarbonyl (Fmoc) protected building block as described earlier [11, 12]. Using this glycosylated Asn building block, a synthetic glycopeptide carrying a homogenous *N*-glycan ($Hex_5HexNAc_4NeuAc_2$) was generated using an established method for solid phase peptide synthesis[12-14]. The synthetic peptide sequence mimicked a tryptic glycopeptide from human vitamin K-dependent protein C present in human serum (UniProtKB, P04070, $^{284}$EVFVHPNYSK$^{293}$). The structure, purity and integrity after deprotection and purification were confirmed using reversed phase LC-MS/MS as described earlier[13].

*Study sample*

Human serum from a commercial source was used for this study (product number #31876, Thermo Fisher Scientific). As a positive control, 52 fmol of the synthetic *N*-glycopeptide from human vitamin K-dependent protein C (see details above), was spiked into 5 µg human serum prior to digestion. Proteins were cysteine reduced and alkylated prior to protein digestion using 1:100 (w/w, enzyme:protein substrate) sequence-grade trypsin for 16 h, 37°C in 20 mM aqueous ammonium bicarbonate, pH 8.0. Undigested protein material and large peptides were removed by filtration using a 30 kDa molecular weight cut off membrane (#88502, Thermo Fisher Scientific). The membrane was washed using 30% (v/v) methanol in 0.1% (v/v) aqueous trifluoroacetic acid (TFA). The flow-through fraction was collected, evaporated using a SpeedVac, and then resuspended in 200 µL 50% (v/v) acetonitrile (ACN) in 0.1% (v/v) aqueous TFA. Glycopeptide enrichment was performed using Hypersep Retain AX columns (#60107-403, Thermo Fisher Scientific). The columns were prepared according to the manufacturer's instructions and were additionally washed with 100 mM aqueous triethylammonium acetate before equilibration with 95% (v/v) ACN in 1% (v/v) aqueous TFA. The sample was diluted in 3 mL 95% (v/v) ACN in 1% (v/v) aqueous TFA, applied to the columns, and then washed with an additional 3 mL 95% (v/v) ACN in 1% (v/v) aqueous TFA before the glycopeptides were eluted with 1 mL 50% (v/v) ACN in 0.5% (v/v) aqueous TFA. The enriched glycopeptide mixtures were dried using a SpeedVac and resuspended in 0.1% (v/v) aqueous TFA for LC-MS/MS analysis.

*Mass spectrometry*

The glycopeptides were separated by reversed-phase nanoLC using a Thermo Scientific EASY-nLC™ 1200 UPLC system connected to a $C_{18}$ LC column (50 cm length × 75 µm inner diameter, Thermo Scientific™ EASY-Spray™). Separation was achieved using a 75 min 6-45% (v/v) and 3 min 45-95 % (v/v) gradient of solvent B consisting of 80% (v/v) ACN in 0.1% (v/v) aqueous formic acid in solvent A consisting of 0.1% (v/v) aqueous formic acid at a 300 nL/min flow rate. The separated glycopeptides were detected using a Thermo Scientific™ Orbitrap Fusion™ Lumos™ Tribrid™ mass spectrometer connected directly to the LC. Approximately 1 µg peptide material was injected on the LC column per run. The same glycopeptide sample was analysed twice using two slightly different acquisition methods producing two related data files (File A and B).

For both methods, MS1 scans were acquired from *m/z* 350 –1,800 in the Orbitrap at a resolution of 120,000 and with an automatic gain control (AGC) of 4 x $10^5$ and an injection time of 50 ms. Data-dependent HCD-MS/MS was performed for the 10 most intense precursor ions selecting the highest charge state and the lowest *m/z* in each MS1 full scan. The HCD-MS/MS fragment ions were recorded in the Orbitrap at a resolution of 30,000 and with an AGC of 5 x $10^4$, injection time of 60 ms, normalised collision energy (NCE) of 28% and a quadrupole isolation width of 2 Th. Already selected precursors were dynamically excluded for 45 s. Product-dependent (pd) ion triggered re-isolation and fragmentation of precursor ions were enabled upon detection of at least one of three selected glycan oxonium ions (*m/z* 138.0545, 204.0867 and 366.1396) if the diagnostic ion(s) was amongst the top 20 fragment ions within each HCD-MS/MS spectrum. For File A, pd-triggered ETciD- and CID-MS/MS events were scheduled. The ETciD-MS/MS fragments were detected in the Orbitrap at a resolution of 60,000 with an AGC of 4 x $10^5$, injection time of 250 ms, CID NCE of 15%, and a quadrupole isolation width of 1.6 Th. Charge-dependent ETD calibration was enabled. The CID-MS/MS fragments were detected in the Orbitrap at a resolution of 30,000 with an AGC of 5 x $10^4$, NCE of 30%, injection time of 54 ms, and a quadrupole isolation width of 1.6 Th. For File B, pd-triggered EThcD- and CID-MS/MS events were scheduled. The EThcD-MS/MS fragments were detected in the Orbitrap at a resolution of 60,000 with an AGC of 4 x $10^5$, injection time of 250 ms, HCD NCE of 15%, and a quadrupole isolation width of 1.6 Th. Charge-dependent ETD calibration was enabled. The CID-MS/MS fragments were detected in the ion trap at unit resolution using a rapid scan method with an AGC of 1 x $10^4$, injection time of 70 ms, NCE of 30%, and a quadrupole isolation width of 1.6 Th. Files A-B were provided to all participants as .raw data files (File A: 684 MB, File B: 811 MB) or as three separate .mgf files containing peak lists of the fragment spectra from the three different fragmentation modes used for File A and B (23.9 MB - 65.6 MB). Conversion to .mgf was performed using ProteoWizard[15].

*Search instructions and reporting template*

The participants were requested to use a protein search space provided by the study organisers comprising the entire human proteome (20,231 UniProtKB reviewed sequences, downloaded January 2018) for their search. In contrast to the fixed protein search space, the participants were free to choose the *N*- and *O*-glycan search space. To limit the number of study variables, participants were asked not to include xylose and any glycan substitutions (e.g. phosphate, sulphate and acetylation) in the glycan search space. The participants were requested to report their team details, identification strategy, and the identified glycopeptides in a common reporting template organised as five separate sheets in an Excel file comprising the following categories of information: 1. Team and contact details, 2. Identification strategy and other study information, 3. *N*- and *O*-glycan search space, 4. List of identified *N*- and *O*-glycopeptides, 5. Summary of identified peptides. The returned reports were carefully checked for compliance with the study guideline. See PXD024101via the PRIDE repository[16] for the common reporting template and the deidentified reports from all participants forming the foundation of this study.

*Search engines and pre- and post-processing tools used for the glycopeptide identification*

A total of 13 search engines were used for glycopeptide identification: IQ-GPA v2.5[1], Protein Prospector v5.20.23[2], glyXtool$^{MS}$ v0.1.4[3], Byonic v2.16.16[4], Sugar Qb[5], Glycopeptide Search v2.0alpha [6], GlycopeptideGraphMS v1.0/Byonic[7], GlycoPAT v2.0[8] and GPQuest v2.0[9], Mascot v2.5.1[17] or v2.2.07, MS Amanda v1.4.14.8243[18], Sequest-HT (in Proteome Discoverer v2.2) (**Extended Data Figure 1h**). These tools were used as stand-alone tools or in combinations with other search engines, while others were applied with pre- or post-processing tools, including OMSSA v2.1.8, Preview v2.13.2, Protein Prospector MS-filter, MS-GF+/PepArML and pParse v.2.0 (**Extended Data Figure 1i**).

*Compilation and comparison of participant reports*

Information of the participating teams was compiled from the returned reports (**Supplementary Table 1-2**). The lists of intact *N*- and *O*-glycopeptides reported by the 22 teams were compiled into a single table with a unique header (**Supplementary Table 3**). Additional columns were manually added to the compiled table with the purpose of standardising some of the reported text variables and generating unique identifiers (IDs) for the reported glycopeptides and their glycan compositions and source glycoproteins. The glycan composition ID was written as the generic monosaccharide composition as Hex*HexNAc*Fuc*NeuAc*, where * represents the number of the individual monosaccharide residues. Glycopeptides adducted with $Na^+$ and $K^+$ were considered and reported by some teams. The adducted glycopeptides were combined with the corresponding non-adducted monosaccharide compositions. UniProtKB identifiers were used as the source protein IDs. The glycopeptide IDs were written as the peptide sequence followed by the generic glycan composition.

The comparisons between the generic glycan compositions, source proteins and glycopeptide IDs reported by the 22 teams were performed using the pivot table tool available in Excel, where the identifier type was placed in "rows", and the team identifier in "columns". The variables from each identifier type were compared as summed counts across the 22 teams.

*Manual quantitative glycoprofiling of select serum N-glycoproteins*

A comprehensive quantitative site-specific analysis of the *N*-glycosylation of four high abundance serum *N*-glycoproteins including alpha-1-antitrypsin (A1AT, UniProtKB, P01009, three *N*-glycosylation sites i.e. Asn70, Asn107 and Asn271), ceruloplasmin (CP, P00450, Asn138, Asn358 and Asn762), haptoglobin (HP, P00738, Asn184 and Asn241) and immunoglobulin G1 (IgG1, P01857, Asn180) was manually performed to allow for a quantitative comparison of the studied glycoprotein sample to glycoprofiling data in the literature[19-22] and thus validate the literature-based performance tests (N2-N3, O1-O2) used to score teams (see below). The site-specific *N*-glycoprofiling data were also used as a "ground truth" to validate the scoring and ranking of teams in an orthogonal manner (see below for details).

For the quantitative site-specific glycoprofiling, the HCD- and EThcD-MS/MS data from File B were firstly searched using Byos v3.9-7 (Protein Metrics Inc., CA, USA)[23, 24]. The "default" search strategy for *N*-glycopeptides commonly used by teams in this study was employed for the Byos search (see details below). The Byos-identified *N*-glycopeptides (PEP-2D < 0.001 was used as a general confidence threshold) were manually confirmed, and the Byos output and the LC-MS/MS raw data were carefully inspected for any additional *N*-glycoforms expected based on the literature of the selected glycoproteins[19-22] or based on biosynthetic rules using Xcalibur v3.0.63 (Thermo Fisher Scientific) and with support from protein sequence handling software GPMAW v9.51 (Lighthouse, Odense, Denmark)[25]. This comprehensive approach ensured that all relevant *N*-glycopeptides belonging to these four source glycoproteins were included in the quantitative analysis. The relative abundance of all observed *N*-glycopeptides from the four selected source glycoproteins was manually determined using EIC-based area-under-the-curve measurements of all observed charge states of the monoisotopic precursor ions using Xcalibur v3.0.63 (Thermo Fisher Scientific). The relative abundance of each glycoform was determined as the percentage of the peak intensity of the individual glycopeptide forms relative to the peak intensity of all glycopeptides spanning each glycosylation site, an approach commonly employed in quantitative glycopeptide analysis[26-28].

*Analysis and optimisation of the search strategies used for the Byonic search engine*

A Byonic-centric analysis and optimisation of the search strategies were performed through a series of controlled in-house searches in which the search settings were systematically varied and the output assessed for performance. For this purpose, only the HCD- and EThcD-MS/MS data from File B were used and searched on an ordinary desktop computer (Windows 10, 64-bit, 16 GB RAM, Intel Core i7-8700 @ 3.20GHz). The "Heavy" multicore parameter option was selected for all searches. Fragment spectra from these two dissociation methods were searched in concert ("HCD/EThcD" setting enabled) using Byonic v3.9.4 (Protein Metrics Inc., CA, USA) using a series of search strategies in which the diverse search settings were sequentially changed. The search strategy used by most teams employing Byonic in this study is herein referred to as the "default" search strategy. The default search strategy employed a predefined glycan database containing either 309 mammalian *N*-glycans or 78 mammalian *O*-glycans available within Byonic, allowed up to one glycan per peptide as a "rare" variable modification, considered only peptides with tryptic cleavage patterns with a maximum of two missed tryptic cleavages per peptide, allowed up to 10/20 ppm deviation of the observed precursor/product ion masses from the expected values, considered up to one Met oxidation (+15.994 Da) per peptide (variable "common" modification), used monoisotopic correction (error check = +/- floor (mass in Da / 4000)), and employed a decoy and contaminant database available in Byonic. One or more of the search settings (SS1-SS2, SS6-SS14) used for the default search strategy were then systematically changed; these alternative settings were selected based on literature and by taking inspiration from search strategies used by the high-performance teams. For the *N*- and *O*-glycan databases (SS1-SS2), customised glycan databases of 25 *N*-glycans expected in human serum (Clerc et al.)[29] or 13 *O*-glycans expected in human serum (Yabu et al.)[30] were used. The systematic searches also explored the output when allowing up to two glycans per peptide as a "rare" variable modification (SS9), when considering semi-specific trypsin cleavages (SS6) with a maximum of one missed cleavage per peptide (SS7), when allowing 5/10 ppm deviation of the observed precursor/product ion masses to their expected values (SS11/SS12), when considering up to four variable "common 1" modifications per peptide, including Met oxidation (+15.994 Da), Asn/Gln deamidation (+0.9840 Da), Gln → pyro Glu (-17.0265 Da) (SS8 and SS10), and, finally, also when no error check for monoisotopic correction (SS14) and no decoy/contaminant database (SS13) were employed. Cys carbamidomethylation (+57.021 Da) (fixed modification) and the protein search space (20,201 entries, all reviewed UniProtKB human proteins, downloaded July 2017) remained constant across all searches and none of the searches employed spectral recalibration. Glycopeptides were filtered to 0% FDR at the peptide level by manually removing glycopeptides identified in the decoy or contaminant database after a general confidence score threshold was applied to the data output (Byonic score >100). The resulting lists of glycopeptides identified from each of these Byonic-centric searches were subjected to the devised performance tests for *N*- and *O*-glycopeptides (N1-N6 and O1-O5, respectively) and the relative sensitivity and specificity scores were determined as described below. All sensitivity and specificity scores were normalised to the scores arising from the default search strategy (set to 1). A detailed summary of the search settings and performance scores generated from these systematic searches can be found in **Supplementary Table 19a**.

In addition to the analysis and optimisation of the search settings used for Byonic, we analysed the impact of the data input on the performance of this search engine. For this purpose, series of controlled searches were carried out by systematically changing the fragmentation type considered for the searches while keeping the search settings and data output filtering constant. The File B raw data file was used as input for all searches and the fragmentation type for each search was specified within the Byonic interface. The "default" search strategy for *N*-glycopeptides was used for all searches (see above for details). Fragment mass tolerance of 0.5 Da and 20 ppm were considered for CID- and HCD/EThcD-MS/MS data, respectively. The following fragmentation types and combinations thereof were tested in individual searches: CID only, HCD only, EThcD only, HCD/EThcD (in concert), HCD/CID (in concert) and HCD/EThcD/CID (in concert). Glycopeptides were filtered using the same criteria described above. Sensitivity and specificity scores were determined for the identified *N*-glycopeptides from each of the searches. Data from these additional searches can be found in **Supplementary Table 19c**.

*Performance testing of teams and software*

The relative team and software performance for glycopeptide data analysis was in this study determined via three different methods as detailed below. In short, all teams were firstly scored and ranked based on a comprehensive assessment method involving multiple complementary performance tests (i). Subsequently, the scoring of teams was validated using an independent glycoprotein-based assessment score (ii). Finally, for the search engine-centric analysis and optimisation of the search strategies for Byonic, the relative performance was evaluated based on a scoring method that produced relative specificity and sensitive scores (iii).

i) Scoring and ranking of teams via multiple performance tests (N1-N6, O1-O5)

The relative team performance was assessed using a scoring system composed of multiple independent tests designed to score the accuracy (specificity) and coverage (sensitivity) of the reported *N*- and *O*-glycopeptides in orthogonal ways. The raw scores from the individual tests (N1-N6 and O1-O5, described below) were normalised within the range 0-1. These normalised scores were used to establish an overall performance score (range 0-1), measuring the ability to perform accurate and comprehensive *N*- and *O*-glycopeptide analysis. The overall performance score was utilised to separately rank the developer and expert user teams.

a) The synthetic *N*-glycopeptide test (N1): All MS/MS spectra corresponding to the synthetic *N*-glycopeptide from human vitamin K-dependent protein C (peptide sequence: EVFVHPNYSK, glycan composition: $HexNAc_4Hex_5NeuAc_2$) were manually retrieved and annotated from File A and B. In total, nine MS/MS spectra corresponded to the non-adducted synthetic *N*-glycopeptide in charge state 3+ and 4+ spanning the four applied fragmentation modes (HCD-, ETciD, EThcD- and CID-MS/MS) (**Extended Data Figure 8a-b**). Further three MS/MS spectra (HCD-, EThcD- and CID-MS/MS) corresponded to the $K^+$-adducted synthetic *N*-glycopeptide in charge state 5+. The sensitivity of the test was determined as the proportion of the 12 MS/MS spectra mapping to the synthetic *N*-glycopeptide that was reported by each team adjusting for the type of fragmentation mode(s) included in their respective search strategies. The specificity was calculated by the proportion of correctly reported glycoPSMs corresponding to the synthetic glycopeptide that matched the 12 annotated MS/MS spectra, again adjusting for the type of fragmentation mode(s) included in the applied search strategies. The test score was calculated by multiplying the sensitivity and specificity (**Extended Data Figure 8c-d** and **Supplementary Table 5**).

b) The glycan composition test (N2 and O1). The *N*-glycan composition score was calculated based on the Pearson correlation ($R^2$) between the expected distribution of *N*-glycans carried by human serum glycoproteins as reported by Clerc et al.[29] and the observed *N*-glycan distribution reported by each team. The *O*-glycan composition score was calculated based on the Pearson correlation ($R^2$) between the expected distribution of *O*-glycans carried by human serum glycoproteins as reported by Yabu et al.[30] and the observed *O*-glycan distribution reported by each team. The distribution of the *N*- and *O*-glycan compositions was calculated based on the glycoPSM count of each unique glycan ID relative to the total glycoPSM count reported by each team.

c) The source glycoprotein test (N3 and O2). The source glycoprotein score was determined from the accuracy (specificity) and coverage (sensitivity) of the reported source glycoproteins relative to the glycoproteins expected in human serum. Reported *N*-glycoproteins previously identified in human serum by both Clerc et al.[29] and Sun et al.[31] received a score of 2, whereas *N*-glycoproteins only identified by Sun et al. received a score of 1. Source glycoproteins not identified by any of the two studies received no score. Further, reported *O*-glycoproteins previously identified in human serum by Darula et al.[32], Yang et al.[33] and Ye et al.[34] received a score of 3, 2 or 1 according to the number of papers identifying the specific *O*-glycoprotein. The source glycoproteins not reported by any of these three studies received no score. For both the serum *N*- and *O*-glycoproteins, the number of glycoPSMs reported by each team was multiplied by the respective source glycoprotein score for each unique glycoprotein ID. The specificity of the test was calculated based on the summed glycoprotein score divided by the highest possible total score (number of unique glycoproteins reported by each team multiplied by the highest theoretical glycoprotein score). The sensitivity of the test was calculated based on the summed number of glycoproteins with score > 0 divided by the number of unique source glycoproteins reported in the selected literature.

d) The glycoproteome coverage test (N4 and O3): The *N*- and *O*-glycoproteome coverage was calculated based on the number of unique glycopeptide IDs (unique peptide sequence and glycan composition) reported by each team.

e) The commonly reported ('consensus') glycopeptide test (N5 and O4): The consensus *N*-glycopeptide score was calculated based on the proportion of glycopeptide ID commonly reported by at least 50% of the 22 teams returning *N*-glycopeptide data. The consensus *O*-glycopeptide score was calculated based on the number of glycopeptide ID commonly reported by at least 30% of the 20 teams returning *O*-glycopeptide data.

f) The NeuGc and multi-Fuc glycopeptide test (N6 and O5). The number of reported *N*- and *O*-glycoPSMs corresponding to NeuGc and multi-Fuc (Fuc ≥ 2) containing glycopeptides was normalised to the total glycoPSMs reported by each team. Separate *N*- and *O*-glycopeptide scores were then calculated based on the average of non-NeuGc and non-Fuc ≥ 2 containing glycoPSMs for teams that included NeuGc and Fuc ≥ 2 containing glycan compositions in their glycan search space.

The overall performance scores for *N*- and *O*-glycopeptide analysis were established separately by averaging the scores of the individual performance tests (N1-N6 and O1-O5, respectively).

ii) Orthogonal glycoprotein-based scoring to validate the team scoring and ranking

To validate the scoring and ranking of teams based on the multiple performance tasks described above (i), an orthogonal glycoprotein-centric scoring method was devised. The method, founded on a "ground truth" as opposed to inference from literature, evaluated the quantitative match of the glycoPSMs reported by the teams to the actual site-specific *N*-glycosylation of selected high-abundance glycoproteins including A1AT, CP, HP and IgG1. For this purpose, two metrics were developed (specificity and sensitivity) to score the match to the actual site-specific glycoform distribution. First, the site-specific distribution of *N*-glycans covering Asn70, Asn107 and Asn271 from A1T1, Asn138, Asn358 and Asn762 from CP, Asn184 and Asn241 from HP and Asn180 from IgG1 was manually determined using AUC-based glycopeptide quantitation (see above for details) and also determined for each team based on spectral counting of reported glycoPSMs. Specificity score was then calculated by multiplying the site-specific glycoform distributions reported by teams by the relative abundance of the actual site-specific glycoforms. The site-glycoform specificity scores were summed within each protein and normalised across the teams (best coverage set to 1). The "overall specificity score" was calculated by averaging the normalised scores from A1AT, CP, HP and IgG1. Sensitivity score was calculated by the proportion of reported non-redundant (unique) glycoforms covering the expected site-specific glycoforms of the four glycoproteins based on robust literature[19-22, 29]. The site-glycoform sensitivity scores were summed within each protein and normalised across the teams (best coverage set to 1). The "overall sensitivity score" was determined by averaging the normalised scores from A1AT, CP, HP and IgG1. Combined scores ("Glycoprotein-centric score") were established by averaging the overall specificity scores and the overall sensitivity scores. The combined scores were then compared to the overall *N*-glycopeptide scores generated from the performance tests N1-N6 (i, see above) using

Pearson correlation ($R^2$). The data underpinning this scoring method can be found in **Supplementary Table 17**.

iii) Search engine-centric scoring of the sensitivity and specificity of Byonic search strategies

To establish the performance of various Byonic search strategies, a scoring method that assessed the relative sensitivity (coverage) and specificity (accuracy) of the search engine was devised. For this purpose, the multiple performance tests already established for the scoring and ranking of teams (N1-N6, O1-O5, see i) were used, but in a slightly different manner. The individual sensitivity and specificity scores from the synthetic glycopeptide test (N1) and source glycoprotein test (N3 and O2) were namely separately considered and grouped with several sensitivity- and specificity-centric performance tests to establish "global sensitivity scores" and "global specificity scores" that could be compared between searches. The global sensitivity score for *N*-glycopeptides was determined by averaging the normalised sensitivity scores from the synthetic glycopeptide (N1), source *N*-glycoprotein (N3), glycopeptide coverage (N4) and commonly reported ('consensus') *N*-glycopeptide (N5) tests. The global specificity for *N*-glycopeptides was determined by averaging the normalised specificity score from the synthetic glycopeptide (N1), glycan composition (N2), source *N*-glycoprotein (N3) and non-NeuGc/multi-Fuc glycopeptide (N6) tests. The global sensitivity score for *O*-glycopeptides was determined by averaging the normalised sensitivity score from source *O*-glycoprotein (O2), glycopeptide coverage (O3) and commonly reported ('consensus') *O*-glycopeptide (O4) tests. The overall specificity for *O*-glycopeptides was determined by averaging the normalised specificity score from the glycan composition (O1), source *O*-glycoprotein (O2) and non-NeuGc/multi-Fuc glycopeptide (O5) tests.

*Statistical analysis*

The normalised performance scores from each performance test were compiled with the search parameters and search outputs (average of selected variables). Seven statistical methods were applied to identify search settings and search output characteristics that were associated with high performance scores including 1) a multiple linear regression model applied with a significance threshold of $p < 0.05$ to identify association between search variables (predictors) and performances scores (response variable), 2) a ridge linear regression model applied using an induced smoothing paradigm for hypothesis testing[35, 36], 3) a Lasso linear model for variable selection[37], 4) a least angle regression exploiting exact post-selection inference to identify associations[38, 39], 5) a forward stepwise linear regression applied using selective inference to identify association[40], 6) a Random Forest algorithm (an ensemble learning model for regression) applied using a variable of importance score to identify association[41] (a permutation strategy on augmented set of noise variables was exploited to define the variable importance cut-off), and 7) a gradient boosting tree algorithm (an ensemble of decision trees for prediction) applied using a similar strategy as the Random Forest algorithm to select important associations[42, 43]. R package v1.2, 1.2.5 and 2.1.8 were used for these analyses. Only associations commonly observed across a minimum of three different statistical methods were considered in this study.

Unpaired two-sided t-tests were applied to compare *N*-glycopeptide (n = 22) against *O*-glycopeptide (n = 20) search output data from all teams (**Extended Data Figure 3**) and to compare the performance scores based on HCD-MS/MS data (n = 17 or n = 16) against EThcD-MS/MS data (n = 13 or n = 10) (**Extended Data Figure 9**). The confidence interval was set to 95% and statistical significance was indicated as $*p < 0.05$, $**p < 0.01$ and $***p < 0.001$.

Pearson correlations ($R^2$) were used to determine i) the quantitative match between the observed site-specific *N*-glycan distribution of four selected glycoproteins in the investigated sample and the site-specific glycoform distribution reported by the literature and ii) the similarity between the overall team scores and the glycoprotein-based scores for all 22 teams.

### References used in the Supplementary Information

38. Hastie, T. & Efron, B. Lars: Least Angle Regression, Lasso and Forward Stagewise. R package version 1.2. (2013).

39. Tibshirani, R.J., Jonathan, T., Lockhart, R. & Tibshirani, R. Exact post-selection inference for sequential regression procedures. arXiv:1401.3889. (2014).

40. Tibshirani, R. et al. SelectiveInference: Tools for Post-Selection Inference. R package version 1.2.5. (2019).

41. Breiman, L. Bagging Predictors. *Machine Learning* **24**, 123–140 (1996).

42. Efron, B. & Hastie, T. Computer Age Statistical Inference: Algorithms, Evidence, and Data Science. (Cambridge University Press, 2016).

43. Greenwell, B., Boehmke, B. & Cunningham, J. (R package version 2.1.8; 2020).
